# Calsyntenin-3 directly interacts with neurexins to orchestrate excitatory synapse development in the hippocampus

**DOI:** 10.1101/2020.01.24.918722

**Authors:** Hyeonho Kim, Dongwook Kim, Jinhu Kim, Hee-Yoon Lee, Dongseok Park, Hyeyeon Kang, Keiko Matsuda, Fredrik H. Sterky, Michisuke Yuzaki, Jin Young Kim, Se-Young Choi, Jaewon Ko, Ji Won Um

## Abstract

Calsyntenin-3 (Clstn3) is a postsynaptic adhesion molecule that induces presynaptic differentiation via presynaptic neurexins (Nrxns), but whether Nrxns directly bind to Clstn3 has been a matter of debate. Here, we show that β-Nrxns directly interact via their LNS domain with Clstn3 and Clstn3 cadherin domains. Expression of splice site 4 (SS4) insert-positive β-Nrxn variants, but not insert-negative variants, reversed the impaired Clstn3 synaptogenic activity observed in Nrxn-deficient neurons. Consistently, Clstn3 selectively formed complexes with SS4-positive Nrxns *in vivo*. Neuron-specific Clstn3 deletion caused significant reductions in number of excitatory synaptic inputs, and moderate impairment of light-induced anxiety-like behaviors in mice. Moreover, expression of Clstn3 cadherin domains in CA1 neurons of Clstn3 conditional knockout mice rescued structural deficits in excitatory synapses, especially within the stratum radiatum layer. Collectively, our results suggest that Clstn3 links to SS4-positive Nrxns to induce presynaptic differentiation and orchestrate excitatory synapse development in specific hippocampal neural circuits.

## Introduction

Synaptogenic adhesion molecules, a class of synaptic transmembrane proteins that induce synaptic differentiation *in vitro* (Missler et al., 2012; Südhof, 2017; Um and Ko, 2013), are central to various aspects of synapse development, but their precise roles in synapse assembly, validation, and/or plasticity *in vivo* are only beginning to be revealed (Ko et al., 2015; Missler et al., 2012; Südhof, 2018). Presynaptic neurexins (Nrxns) and leukocyte common antigen-related receptor protein tyrosine phosphatases (LAR-RPTPs) are among the synaptogenic adhesion molecules that have emerged as key platforms that facilitate convergence of diverse signals from multifarious postsynaptogenic ligands at mammalian synapses (Han et al., 2016; Südhof, 2018). Of particular note is the fact that, although Nrxns and LAR-RPTPs are evolutionarily conserved, only a subset of their ligands in mammals has homologs in invertebrate species that also play crucial roles in various aspects of central nervous system development, hinting at the possibility that Nrxns and LAR-RPTPs serve fundamental functions through these selective adhesion pathways.

Calsyntenins (Clstns) are evolutionarily conserved synaptogenic adhesion proteins of the cadherin superfamily that are expressed most highly in the brain (Sotomayor et al., 2014). Synaptic functions of the three vertebrate Clstn family members have recently been reported. For example, juvenile Clstn1-deficient mice exhibit compromised excitatory synaptic transmission, possibly owing to disrupted targeting of N-methyl-D-aspartate (NMDA) receptor subunits (Ster et al., 2014). They also show increased synaptic levels of GluN2B subunit-containing NMDA receptors, enhanced long-term potentiation (LTP) and greater filopodia-like dendritic protrusions in the hippocampus, but decreased dendritic arborization, suggesting that Clstn1 mediates dendritic transport of NMDA receptor subunits and regulates spine maturation during early development (Ster et al., 2014). In addition, Clstn1 regulates guidance receptor trafficking by shuttling Rab11-positive vesicles, leading to switching of commissural axon responsiveness (Alther et al., 2016). Clstn1 also contributes to peripheral sensory axon arborization, branching, endosomal dynamics, and microtubule polarity in zebrafish (Lee et al., 2017; Ponomareva et al., 2014). Clstn2, on the other hand, plays a non-redundant role in inhibitory synapse development and influences a subset of cognitive abilities (Lipina et al., 2016; Ranneva et al., 2017). Clstn3 was identified as a postsynaptogenic adhesion molecule that acts through presynaptic Nrxns (Pettem et al., 2013; Um et al., 2014). However, in contrast to the report of Pettem *et al*. (2013), we did not detect direct interactions between Clstn3 and α-Nrxns.

In the present study, we revisited these issues. Strikingly, utilizing newly engineered Nrxn1 expression vectors to increase Nrxn1β expression levels, we found that recombinant Clstn3 bound both Nrxn1β and Nrxn1α, with a slight preference for splice site 4 (SS4) insert-positive variants, requiring an Nrxn splice variant containing an insert at SS4 as a functional receptor for its presynaptic differentiation-inducing activity. Conditional deletion of Clstn3 in neurons led to drastic reductions in excitatory synapse structures (but not basal excitatory synaptic transmission), and mild impairments in a subset of hippocampus-dependent mouse behaviors. Finally, adeno-associated virus-mediated expression of Nrxn-binding CST-3 cadherin domains was sufficient to rescue decreased excitatory synapse puncta density in CA1 stratum radiatum layers of Clstn3-deficient mice. Viewed together, our results revise our previous molecular model, showing that Clstn3 directly interacts with SS4-positive Nrxn splice variants to induce presynaptic differentiation, and suggesting synapse-organizing functions of Clstn3 that may control specific synaptic inputs from Schaffer-collateral afferents in the hippocampus.

## Results

### Clstn3 directly binds β-Nrxns

Our previous cell surface-binding assays used a variety of Nrxn1α deletion variants derived from the bovine *Nrxn1α* gene or Nrxn1β variants derived from the rat *Nrxn1β* gene (Um et al., 2014). These vectors have long been used to characterize Nrxn interactions with neuroligins (NLs) and other ligands, including leucine-rich repeat transmembrane neuronal proteins (LRRTMs), neurexophilins, and latrophilin-1 (Boucard et al., 2012; Ko et al., 2009a; Missler et al., 1998). However, using Nrxn vectors constructed from mouse *Nrxn* genes, Craig and colleagues reported that Nrxn1α, but not Nrxn1β, binds to Clstn3 (Pettem et al., 2013). To resolve this discrepancy, we performed affinity chromatography of solubilized mouse synaptosomes using immobilized recombinant Ig-Clstn3 followed by mass spectrometry (MS). Intriguingly, among the captured proteins was a tryptic peptide unique to Nrxn1β (in addition to Nrxn1α peptides) (***Figures 1A–D***; see ***Table 1***). To confirm binding between Clstn3 and Nrxn1β, we engineered mouse Nrxn1β expression constructs and performed cell surface-binding assays. We found robust binding of Ig-Clstn3 to HEK293T cells expressing C-terminally FLAG-tagged mNrxn1β lacking (mNrxn1β^−SS4^-FLAG) or containing (mNrxn1β^+SS4^-FLAG) the SS4 insert (***Figures 1E*** and ***1F***). In assays measuring dimeric ligand binding to mNrxn-expressing cell surfaces, Ig-Clstn3 interacted with both mNrxn1β^-SS4^ and mNrxn1β^+SS4^ with nanomolar affinity (***Figures 1G*** and ***1H***). *Kd* values calculated by Scatchard analyses were 51.39 ± 5.26 nM for Nrxn1β^+SS4^ and 105.94 ± 7.89 nM for Nrxn1β^-SS4^ (***Figures 1G*** and ***1H***). These findings indicate that Clstn3 binds Nrxn1β with high affinity and exhibits a slight preference for SS4 insert-positive splice variants. To investigate differences between bovine and rat Nrxn1 plasmids (bNrxn1α and rNrxn1β) used in our previous studies and the mouse Nrxn1 plasmids (mNrxn1α and mNrxn1β) used in the current study, we compared total protein expression levels produced by the different vectors by immunoblot analysis using serially diluted lysates from HEK293T cells transfected with expression plasmids for mNrxn and bNrxn1α or rNrxn1β (***Figure 1—figure supplement 1***). Surprisingly, expression levels of mNrxn vectors were ∼100-fold higher than those of bNrxn and rNrxn vectors (***Figure 1—figure supplement 1A***). This difference in expression was not likely attributable to differences in the position of the FLAG epitope between rNrxn1β (N-terminus) and our original mNrxn1β construct, as evidenced by the comparably robust binding of a newly generated N-terminally FLAG-tagged mNrxn1β^+SS4^ (FLAG-mNrxn1β^+SS4^) and our original C-terminally FLAG-tagged mNrxn1β^+SS4^ (mNrxn1β^+SS4^-FLAG) to Ig-Clstn3 in cell surface-binding assays (***Figure 1—figure supplement 1B and 1C***).

**Figure 1.**
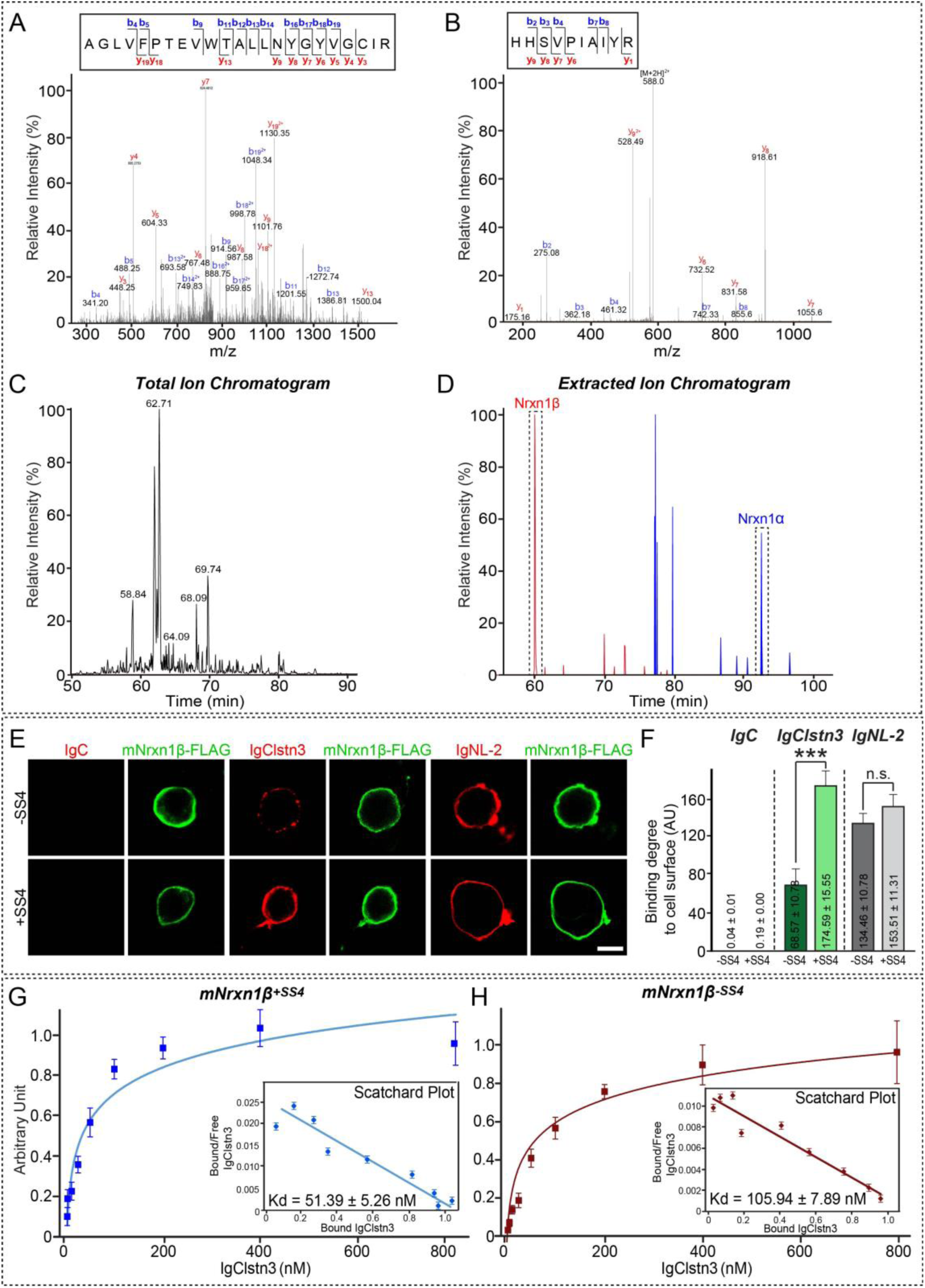
Analysis of Clstn3–β-Nrxn interactions by MS and cell surface-binding assays. **(A** and **B)** MS/MS spectra of two double-charged peptides unique to Nrxn1α and Nrxn1β at m/z 867.11 and 596.33, respectively, were obtained by liquid chromatography (LC)-MS/MS and were fragmented to produce MS/MS spectra with b- and y-ion series describing the sequences AGLVFPTEVWTALLNYGYVGCIR (amino acids [aa] 501–523) and HHSVPIAIYR (aa 64–73). **(C** and **D)** MS data. (**C**) Total ion chromatogram (XIC) of tryptic digests of Ig-Clstn3–bound eluates separated by LC. (**D**) Extracted ion chromatograms of m/z 867.11 and 596.33 ions from Nrxn1α (92.8 min) and Nrxn1β (60.1 min), respectively. **(E)** Cell surface-binding assays. HEK293T cells expressing C-terminally FLAG-tagged mouse Nrxns were incubated with purified Ig-fused neuroligin-2 (Ig-NL-2), calsyntenin-3 (Ig-Clstn3), or negative control (Ig-C) and analyzed by immunofluorescence imaging for Ig-fusion proteins (red) and FLAG (green). Scale bars: 10 μm (applies to all images). **(F)** Quantification of cell surface binding in **(E)**. Data are means ± SEMs (\*\*\**p* < 0.001; non-parametric Kruskal-Wallis test with Dunn’s *post hoc* test). **(G** and **H)** Saturation binding of Ig-Clstn3 to FLAG-tagged mouse Nrxn1β^+SS4^ (**G**) or Nrxn1β^-SS4^ (**H**) expressed in HEK293T cells. Inset: Scatchard plot generated by linear regression of the data; K_d_ was calculated from three independent experiments. Data are means ± SEMs.

### Cadherin domains of Clstn3 mediate direct binding to β-Nrxns

In addition to cell surface–binding assays*, in vitro* pull-down assays clearly showed binding of Clstn3 to both Ig-mNrxn1β^+SS4^ and Ig-mNrxn1α^+SS4^. To identify a minimal Clstn3 domain involved in Nrxn binding, we used three different FLAG-tagged Clstn3 constructs: Full (full-length Clstn3), Cad (containing tandem cadherin domains, the transmembrane segment and intracellular residues of Clstn3), and ΔCad (full-length Clstn3 lacking the tandem cadherin domains) (see ***Figure 2A*** for schematic diagram of Clstn3 constructs). Both Ig-mNrxn1α^+SS4^ and Ig-mNrxn1β^+SS4^ pulled down Clstn3 Cad, but not Clstn3 ΔCad (***Figures 2B*** and ***2C***), suggesting that cadherin domains of Clstn3 mediate binding to Nrxns. The results of these and aforementioned cell-based surface-binding assays do not exclude the possibility that intermediate(s) expressed in HEK293T cells may bridge indirect associations of Nrxns with Clstn3. However, binding assays performed using purified recombinant Ig-Nrxn1 (Ig-Nrxn1β^-SS4^, Ig-Nrxn1β^+SS4^, Ig-Nrxn1α^-SS4^, or Ig-Nrxn1α^+SS4^) and recombinant His-HA-Clstn3 (Clstn3 Ecto, Clstn3 Cad, or Clstn3 ΔCad) (***Figure 2D***) showed that His-HA-Clstn3 Ecto directly bound to recombinant α- and β-Nrxns (***Figures 2E*** and ***2F***). Moreover, Clstn3 Cad, but not Clstn3 ΔCad, directly bound to recombinant β-Nrxns (***Figure 2F***). These data suggest that cadherin domains of Clstn3 mediate direct interactions with Nrxns, consistent with our previous observation that the cadherin domains of Clstn3 are necessary and sufficient to induce presynaptic differentiation (Um et al., 2014).

**Figure 2.**
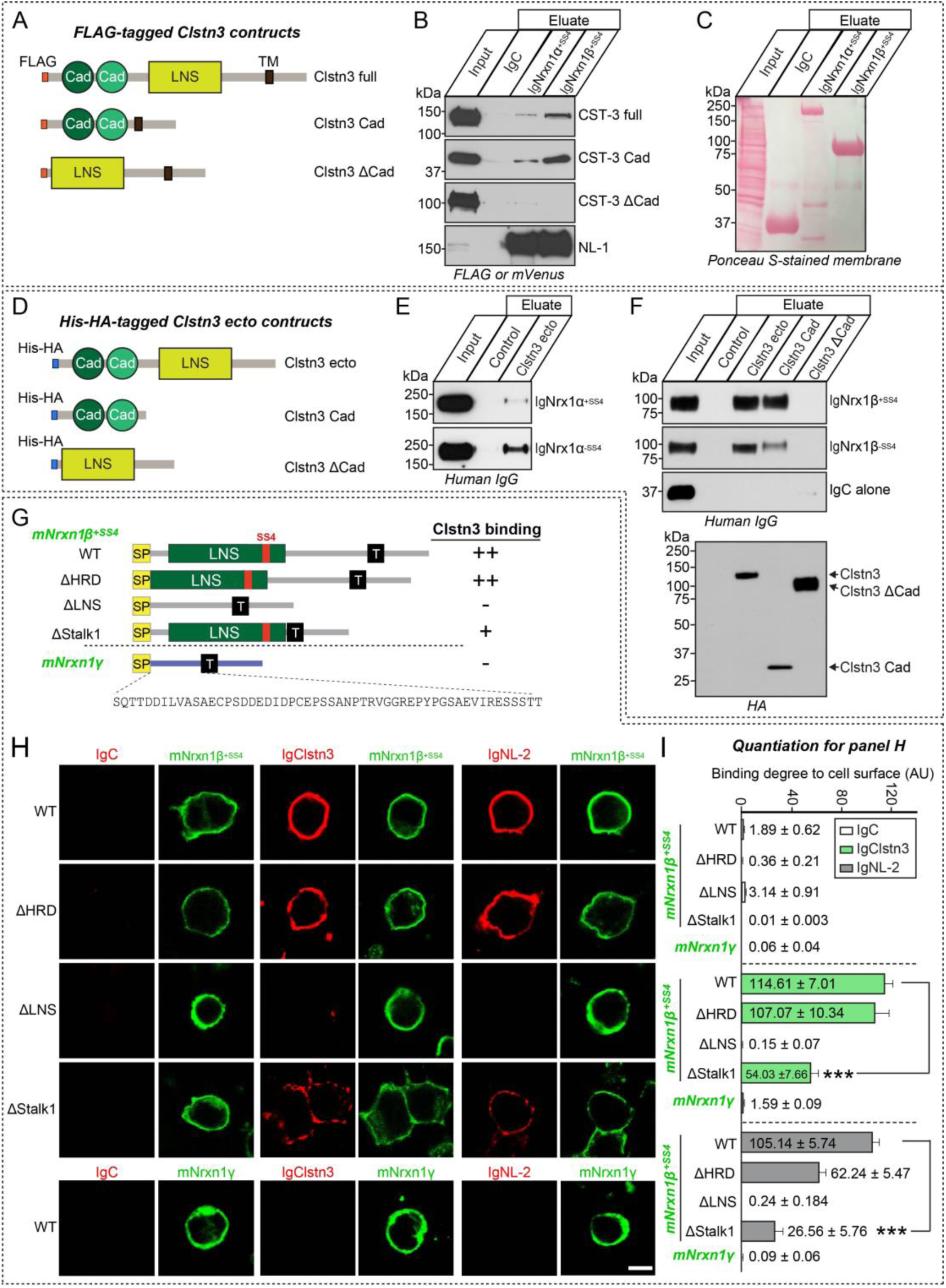
Direct binding of Clstn3 cadherin domains to β-Nrxns. **(A)** Diagrams of Clstn3 variants used in pull-down assays presented in **(B)**. **(B)** Ig-Nrxn1α^+SS4^, Ig-Nrxn1β^+SS4^, or Ig-C (negative control) proteins pulled down FLAG-Clstn3 Full and FLAG-Clstn3 Cad, but not FLAG-Clstn3ΔCad. Input, 5%. **(C)** Ig-Nrxn1α^+SS4^, Ig-Nrxn1β^+SS4^ or Ig-C used for pull-down assays were analyzed by direct comparison of bands revealed by parallel Ponceau S staining. **(D)** Diagrams of the His-HA-tagged extracellular Clstn3 variants used in direct binding assays. **(E) (E** and **F)** Purified His-tagged Clstn3 proteins were incubated with purified Ig-Nrxn proteins, as indicated. Precipitates obtained using Talon resin were analyzed by immunoblotting with human IgG or HA antibodies. Input, 10%. **(G)** Diagrams of Nrxn1β and Nrxn1γ constructs used in **(H)**. **(H)** Cell surface-binding assays. HEK293T cells expressing FLAG-tagged Nrxn1β^+SS4^ WT, its deletion variants (ΔHRD, ΔLNS, or ΔStalk1), or FLAG-tagged Nrxn1γ WT were incubated with Ig-Clstn3, Ig-NL-2 or Ig-C, and analyzed by immunofluorescence imaging for Ig-fusion proteins (red) and FLAG (green). Scale bars: 10 μm (applies to all images). **(I)** Quantification of cell surface binding in **(H)**. Data are means ± SEMs (\*\*\**p* < 0.001; non-parametric Kruskal-Wallis test with Dunn’s *post hoc* test; number of cells analyzed = 11–15).

### Clstn3 binds to an LNS domain of mNrxn1β in a Ca^2+^-dependent manner

We next investigated which Nrxn sequences mediate Clstn3 binding. To this end, we generated the following FLAG-tagged mNrxn1β-deletion constructs: mNrxn1β ΔHRD, which lacks a β-Nrxn-unique histidine-rich sequence; mNrxn1β ΔLNS, which lacks a β-Nrxn LNS domain; and ΔStalk1, which lacks the entire stalk region (***Figure 2G***). We found that Ig-Clstn3 bound comparably to HEK293T cells expressing mNrxn1β ΔHRD or mNrxn1β wild-type (WT), but did not bind HEK293T cells expressing mNrxn1β ΔLNS (***Figures 2H*** and ***2I***), indicating that the Nrxn1β LNS domain is necessary for Clstn3 binding. Interestingly, deletion of the entire stalk region (mNrxn1β ΔStalk1) diminished Clstn3 binding as well as NL-2 binding (***Figures 2H*** and ***2I***). The stalk region contains ∼40 residues with several putative O-linked glycosylated sites and a short cysteine-loop sequence composed of two conserved cysteines flanking an 8-residue acidic sequence (Sterky et al., 2017). We thus hypothesized that O-linked glycosylation of mNrxnβ regulates Clstn3 binding and that mNrxn1β ΔStalk1 displayed weak Clstn3 binding because it lacks O-linked glycosylation. To test this, we generated mNrxn1β constructs containing point mutations or deletions in the conserved stalk region. Clstn3 binding was retained in mNrxn1β constructs in which putative O-glycosylated threonines or serines were replaced with glycines (mNrxn1β ΔCHO), the cysteine-loop sequence was deleted (mNrxn1β ΔCysL), the stalk region was partially deleted (mNrxn1β ΔStalk2), or the conserved serine residue for attaching heparan sulfate chains was replaced with alanine (mNrxn1β ΔHS) (***Figure 2—figure supplement 1A–D***). However, Clstn3 did not bind Nrxn1γ, a newly identified Nrxn1 isoform (Sterky et al., 2017), or other heparan sulfate proteogylcans, such as glypicans or syndecans (***Figure 2H, 2I, Figure 2—figure supplement 1E*** and ***1F***). Control experiments showed that surface expression levels of individual mNrxn1β deletion constructs in HEK293T cells were comparable to those of WT protein (***Figure 2— figure supplement 2***).

These findings suggest that the Nrxn1β LNS domain is a Clstn3-binding site. We further found that treatment with the Ca^2+^ chelator EGTA prevented these interactions (***Figure 2—figure supplement 3***).

### Clstn3 requires specific Nrxn splice variants for presynaptic differentiation

To delineate the molecular mechanisms that link Nrxns to Clstn3 and support its synaptogenic activity, we performed heterologous synapse-formation assays in cultured hippocampal neurons in which all three Nrxns were downregulated using small hairpin RNAs (shRNA) (Um et al., 2016; Um et al., 2014). We found that Nrxns triple-knockdown (TKD) significantly reduced the synaptogenic activity of Clstn3 (***Figures 3A*** and ***3B***) (Um et al., 2014). Synaptogenic activity was completely restored by coexpression of Nrxn1α^+SS4^ or Nrxn1β^+SS4^, but not Nrxn1α^-SS4^ or Nrxn1β^-SS4^, suggesting that SS4-positive Nrxns act as functional receptors for Clstn3 (***Figures 3A, 3B*, *Figure 3—figure supplement 1***). Similarly, re-expression of the respective Nrx splice variants rescued deficits in the synaptogenic activities of NL-1 (neuroligin-1) and LRRTM2 (leucine rich repeat transmembrane neuronal 2) (***Figures 3A*** and ***3B***). Subsequent pull-down assays showed that both Ig-mNrxn1α^+SS4^ and Ig-mNrxn1β^+SS4^, but not Ig-mNrxn1α^-SS4^ or Ig-mNrxn1β^-SS4^, were capable of capturing Clstn3 from mouse synaptosomal membrane fractions (***Figures 3C*** and ***3D***).

**Figure 3.**
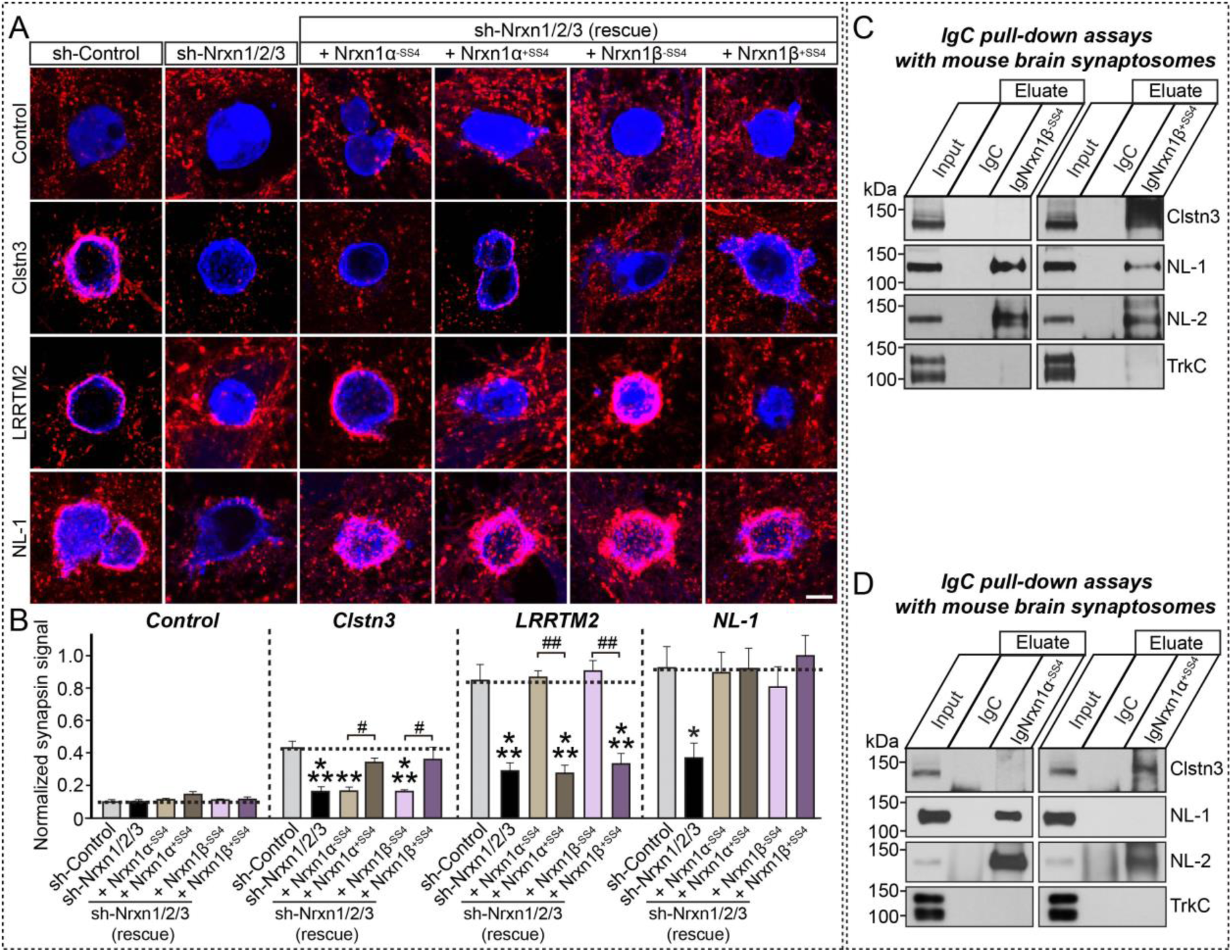
Selected Nrxn variants are required for Clstn3-mediated induction of presynaptic differentiation. **(A and B)** Effects of Nrxn-TKD (sh-Nrnx1/2/3) on the synaptogenic activities of Clstn3, LRRTM2, and NL-1. HEK293T cells expressing the indicated proteins were cocultured with neurons infected with control lentiviruses (sh-Control) or lentiviruses expressing sh-Nrxn1/2/3, without or with coexpression of the indicated Nrxn1α or Nrxn1β splice variants. Representative images **(A)** of cocultures immunostained with antibodies to mVenus or HA (blue) and synapsin I (red). Scale bar: 10 μm (applies to all images). Quantitation **(B)** of heterologous synapse-formation assays, determined by calculating the ratio of synapsin to EGFP/HA fluorescence signals. Dashed lines correspond to control values used as a baseline. Data are means ± SEMs (\*\*\**p* < 0.001; \*\**p* < 0.01; \**p* < 0.05; ^##^*p* < 0.01; and ^#^*p* < 0.05; non-parametric Kruskal-Wallis test with Dunn’s *post hoc* test; ‘n’ denotes the total number of HEK293T cells analyzed). **(C** and **D)** Pull-down assays in solubilized mouse synaptosomal fractions. Assays were performed using recombinant Ig-Nrxn1β (**C**), Ig-Nrxn1α (**D**), or Ig-C proteins. Equal amounts of bound proteins were analyzed using the antibodies indicated to the right of panels. Input, 5%.

### Generation and Characterization of Clstn3 Conditional Knockout (cKO) Mice

To elucidate the physiological significance of Clstn3–Nrxn interactions *in vivo*, we used *Clstn3^tm1a(EUCOMM)Hmgu^* mice in which a targeting cassette harboring FRT, lacZ, and loxP sites was inserted between exon 7 and exon 8, resulting in a ‘knockout-first’ lacZ-reporter-tagged *Clstn3^tm1a^* insertion allele with conditional potential (***Figure 4—figure supplement 1A***) (Skarnes et al., 2011). The *Clstn3^tm1a^* allele was confirmed by genomic polymerase chain reaction (PCR) (***Figure 4—figure supplement 1B***), and loss of Clstn3 protein in *Clstn3^tm1a/tm1a^* mice was verified by the absence of detectable Clstn3 immunoreactivity to two Clstn3-specific antibodies (JK001 and JK091; ***Figure 4—figure supplement 1C*** and ***1D***). Deletion of Clstn3 in *Clstn3^tm1a/tm1a^* mice did not affect the expression levels of diverse synaptic proteins, including Clstn1 and Clstn2 (***Figure 4—figure supplement 1E*** and ***1F***). In addition, the gross morphology of *Clstn3^tm1a/tm1a^* mice was normal and their neuron numbers were comparable to those in WT mice, as assessed by Nissl and NeuN (neuronal marker) staining (***Figure 4—figure supplement 1G–I***). We then crossed *Clstn3^tm1a/tm1a^* mice with an FLPe knock-in strain to remove contaminating transgenes and the neomycin resistance cassette to generate *Clstn3^tm1c(EUCOMM)Hmgu^* mice. Before deciding which Cre driver lines to cross for the current study, we analyzed Clstn3 expression patterns in the mouse brain. For this, we performed RNAscope-based *in situ* hybridization analysis using a *Clstn3*-specific probe. This analysis showed that *Clstn3* mRNA was strongly expressed in most pyramidal cell layers and interneurons in the mouse hippocampus (***Figure 4—figure supplement 2A***). Notably, *Clstn3* expression in pyramidal cell layers in the CA3 region was particularly prominent (***Figure 4—figure supplement 2A*** and ***2C***). *Clstn3* mRNA was also expressed in parvalbumin (PV)- and somatostatin (SST)-positive GABAergic interneurons in the hippocampus, as previously reported (Pettem et al., 2013) (***Figure 4—figure supplement 2B***). Moreover, immunofluorescence analyses using a Clstn3-specific antibody (JK091) showed that the expression pattern of Clstn3 protein was similar to that of *Clstn3* mRNA in the hippocampus (***Figure 4—figure supplement 2D***). Based on these results, the Nestin-Cre (Nestin-*Clstn3*) driver line was chosen to further cross with *Clstn3^tm1c(EUCOMM)Hmgu^* mice to generate Clstn3-cKO mice (***Figures 4A*** and ***4B***). These mice were largely indistinguishable from control littermates (*Clstn3^f/f^*; Ctrl) in terms of birth rate, although Nestin-*Clstn3* mice weighted marginally (but significantly) less at both postnatal day 30 (P30) and P54, which is a generalized metabolic phenotype of the Nestin-Cre driver line (Galichet et al., 2010) (***Figure 4C***; ***Table S2***). In addition, gross morphology (as assessed by Nissl staining) and neuron numbers (as assessed by NeuN staining) in Nestin-*Clstn3* mice were comparable to those of control littermates (***Figure 4D–F***). Semi-quantitative immunoblot analyses showed that the relative expression levels of various synaptic proteins in the hippocampus (***Figure 4G***) and cortex (***Figure 4H***) of Nestin-*Clstn3* mice were unchanged compared with littermate controls (***Figures 4G–I***). Collectively, these data suggest that Clstn3 is not essential for mouse survival or breeding, and does not affect the expression levels of synaptic proteins.

**Figure 4.**
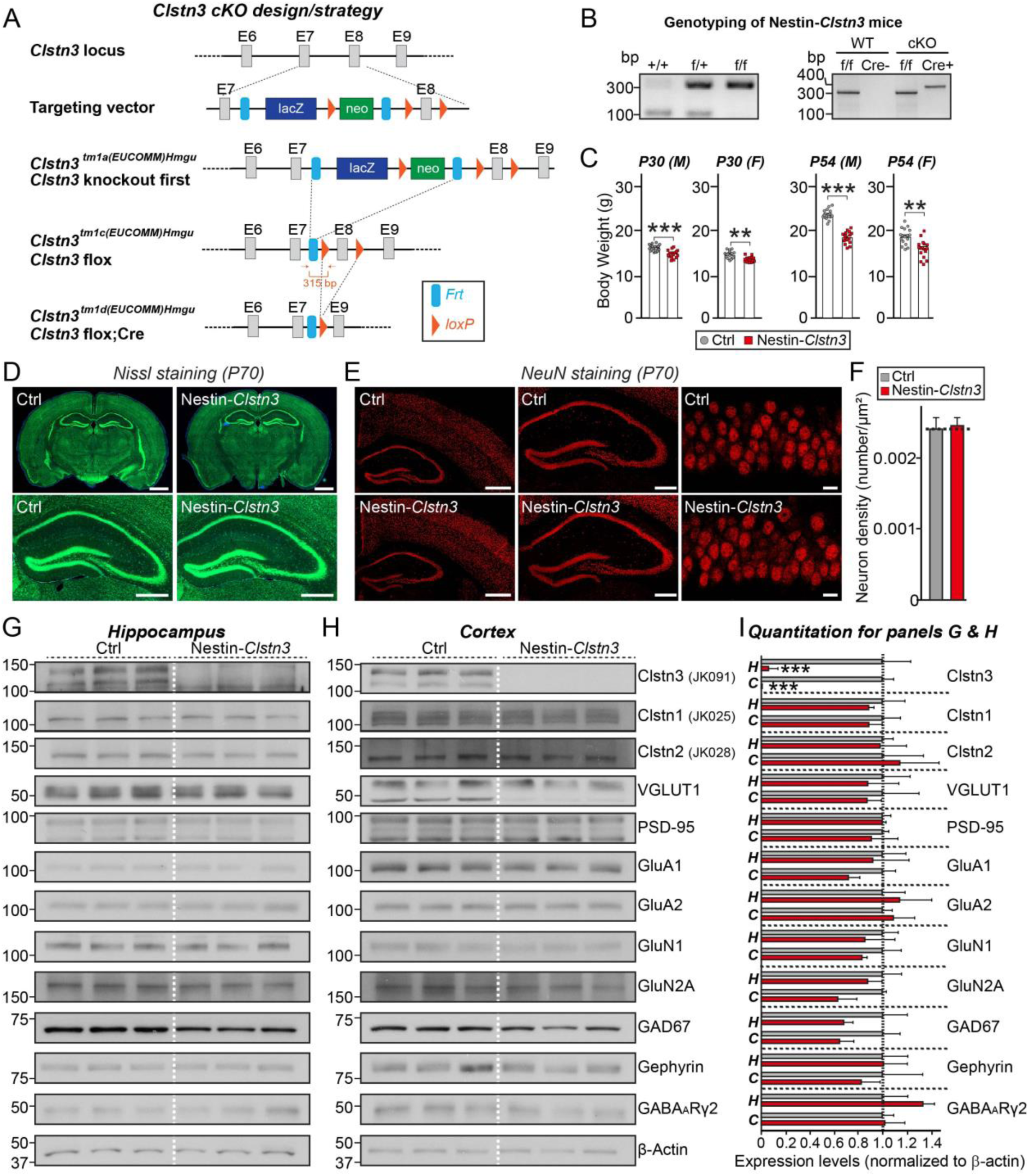
Characterization of Clstn3-cKO mice. **(A)** Strategy used to generate conditional *Clstn3^tm1d/tm1d^* KO mice. Arrows flanking the neomycin gene and FLP recombinant target (FRT) or exon 8 (E8) indicate loxP sites. Red arrows indicate forward and reverse primers used for genotyping. Note that lacZ and neomycin cassettes are two separate markers. **(B)** PCR genotyping of *Nestin-Cre; Clstn3^fl/fl^* (Nestin-*Clstn3;* cKO). The 400 bp Cre-specific PCR product was detected only in Nestin-*Clstn3* mice. The band size of the Clstn3 floxed allele was 315 bp. **(C)** Body weight of Nestin-*Clstn3* and *Clstn3^fl/fl^* (Ctrl) mice at P30 and P54. Abbreviations: F, female; M, male. Data are means ± SEMs (\*\*\**p* < 0.001, \*\**p* < 0.01; Mann-Whitney U test) **(D)** Normal gross morphology of the Nestin-*Clstn3* brain at P70, as revealed by Nissl staining. Scale bar: 1 mm (top) and 500 μm (bottom). **(E)** Representative images of NeuN (a neuronal marker) staining from the Nestin-*Clstn3* brain. Normal numbers of neurons (**F**) in hippocampal and cortical regions (left, forebrain coronal section; middle, hippocampal coronal section; and right, hippocampal CA1 stratum pyramidale layer). Scale bar: 1 mm (left), 0.5 mm (middle), and 20 μm (right). **(F)** Summary graphs of NeuN (neuron marker) staining from (**E**). Data are means ± SEMs (n = 3 mice each after averaging data from 6 sections/mouse; n.s., not significant). **(G–I)** Representative immunoblots for the hippocampus **(G)** and cortex **(H)** and summary graphs of synaptic protein levels (**I**) in crude synaptosomal fractions of P42 Ctrl and Nestin-*Clstn3* brains, analyzed by semi-quantitative immunoblotting. Data are means ± SEMs (\*\*\**p* < 0.001; Mann-Whitney U test; n = 5 mice each group).

### Excitatory synapse development is impaired in CA1 hippocampal pyramidal neurons of Clstn3-cKO mice

We next performed immunohistochemistry to quantify the intensity of excitatory and inhibitory synapses, identified by labeling with antibodies to VGLUT1 (vesicular glutamate transporter 1) and GAD67 (glutamic acid decarboxylase 67), respectively. To this end, we analyzed the effect of Clstn3 deletion on the intensity of excitatory and inhibitory synaptic puncta in various layers of the hippocampal CA1 region (***Figure 5***) using both Nestin-*Clstn3* (**Figure 5**) and *Clstn3^tm1a/tm1a^* mice (***Figure 5—figure supplement 1***). We found a significant decrease in the intensity of VGLUT1 puncta in most layers of hippocampal CA1 regions (stratum oriens [SO] and stratum radiatum [SR]), but not in the stratum lacunosum moleculare (SLM) layer, in both Clstn3-cKO (***Figures 5A*** and ***5C***) and *Clstn3^tm1a/tm1a^* mice (***Figure 5—figure supplement 1***). In contrast, the intensity of GAD67 puncta was unchanged in all examined layers of hippocampal CA1 regions (SO, SR, stratum pyramidale [SP] and SLM) (***Figures 5B, 5D*** and ***Figure 5—figure supplement 1***). To complement these anatomical analyses, we performed whole-cell electrophysiological recordings of miniature excitatory and inhibitory postsynaptic currents (mEPSCs and mIPSCs) in brain slices from Nestin-*Clstn3* and littermate WT mice (***Figure 5—figure supplement 2***). Surprisingly, no differences in the amplitude or frequency of mEPSCs and mIPSCs were detected between Nestin-*Clstn3* and Ctrl mice (***Figure 5—figure supplement 2***). Thus, our data suggest that Clstn3 is specifically required for excitatory synapse structures, but not basal excitatory synaptic transmission, in CA1 hippocampal pyramidal neurons.

**Figure 5.**
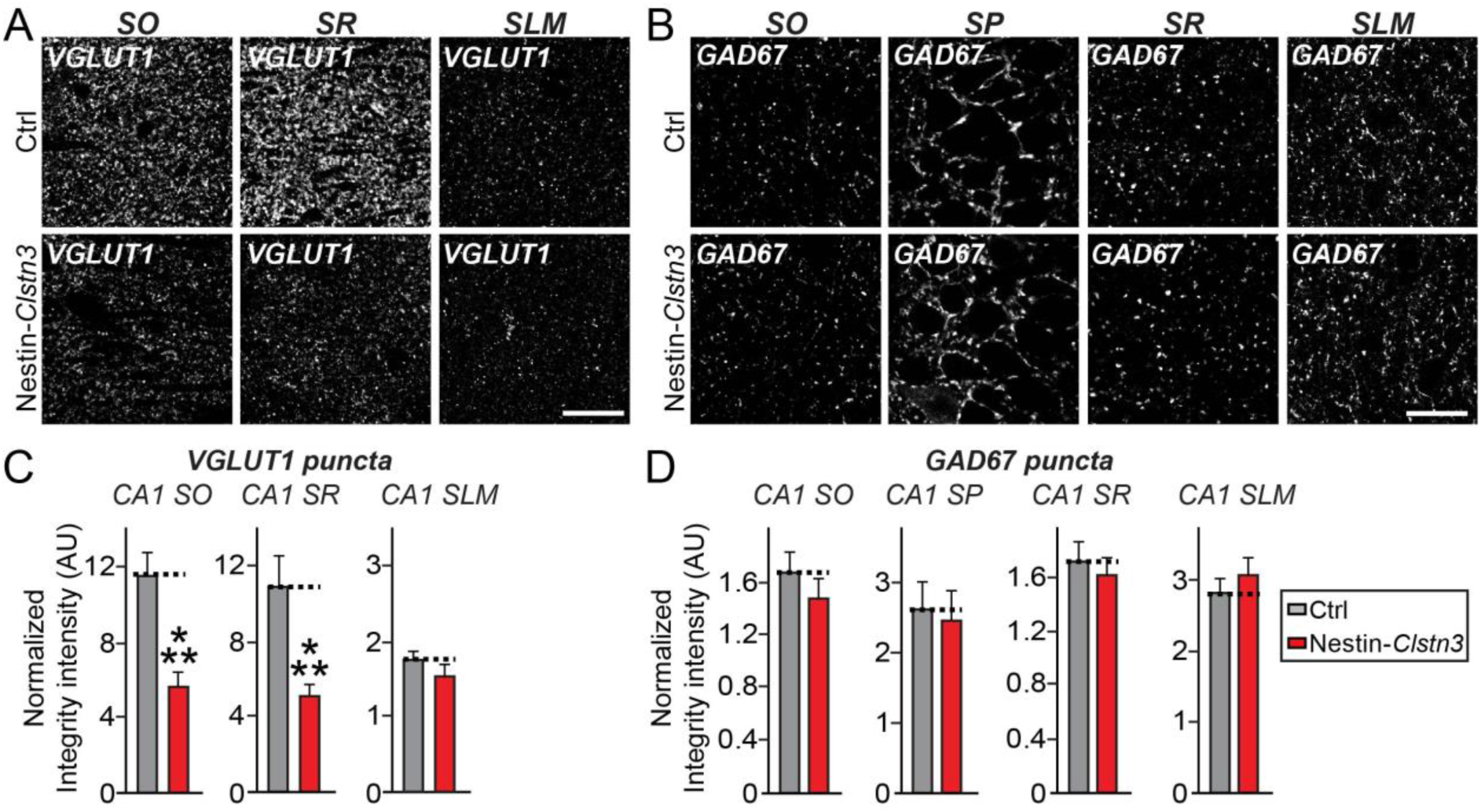
Impaired excitatory synapse development in the hippocampal CA1 region in *Nestin-Cre;Clstn3^fl/fl^* mice. **(A** and **B)** Representative images of hippocampal CA1 SO and SR regions from *Clstn3^fl/fl^* (Ctrl) and *Nestin-Cre;Clstn3^fl/fl^* (Nestin-*Clstn3*) mice, immunostained for the excitatory synapse marker VGLUT1 (**A**) and the inhibitory synapse marker GAD67 (**B**). Scale bar: 20 μm (applies to all images). **(C** and **D)** Quantification of the integrated intensity of VGLUT1-positive synaptic puncta (**C**) from the CA1 SO, CA1 SR and CA1 SLM (left), and GAD67-positive synaptic puncta (**D**) from the CA1 SO, CA1 SP, CA1 SR and CA1 SLM (right). Data are means ± SEMs (\*\*\**p* < 0.001; Mann-Whitney U test; n = 5 mice each group after averaging data from 6 sections/mouse).

### Clstn3 KO mice display increased anxiety-like behavior

To explore the behavioral consequences of Clsnt3 loss in mice, we tested Clstn3-cKO mice for changes in select forms of learning and memory using 8–10-wk-old male Nestin-*Clstn3* mice, together with control male littermates (Ctrl; ***Figure 6***). We initially tested anxiety-like behaviors using the open-field (OF) test, elevated-plus maze (EPM) test, and light/dark t ransition (LDT) test. For the OF test, both Ctrl and Nestin-*Clstn3* mice were allowed to freely explore an open field for 30 min, and then general activity was measured by calculating the velocity and total distance traveled. Anxiety-like behavior was assessed by calculating the frequency of entries into the center of the field and time spent in the center of the field. We found no differences in these parameters in Nestin-*Clstn3* mice compared with control mice (***Figures 6A–E***). There were also no differences between Nestin-*Clstn3* mice and control mice in either the number of entries into open arms or the total time spent in open arms in the EPM test (***Figures 6F–H***). However, in the LDT test, which measures the level of anxiety in response to bright light (unlike the EPM test, which measures the level of anxiety in response to exposure to an open space), there was a significant increase in anxiety-like behavior in Nestin-*Clstn3* mice compared with control mice (both Nestin-Cre and *Clstn3^tm1a/tm1a^* mice; ***data not shown***), measured as the frequency of entering the brightly lit chamber and the total time spent in the lighted chamber (***Figures 6I*** and **6J**). Nestin-*Clstn3* mice displayed normal spatial working memory (measured by the Y-maze test) (***Figures 6K–M***) and short-term memory (assessed by the novel object-recognition (NOR) test) (***Figures 6N–Q***). Overall, these data suggest that Nestin-*Clstn3* mice display moderately increased bright light-induced anxiety, but exhibit normal open space-related anxiety.

**Figure 6.**
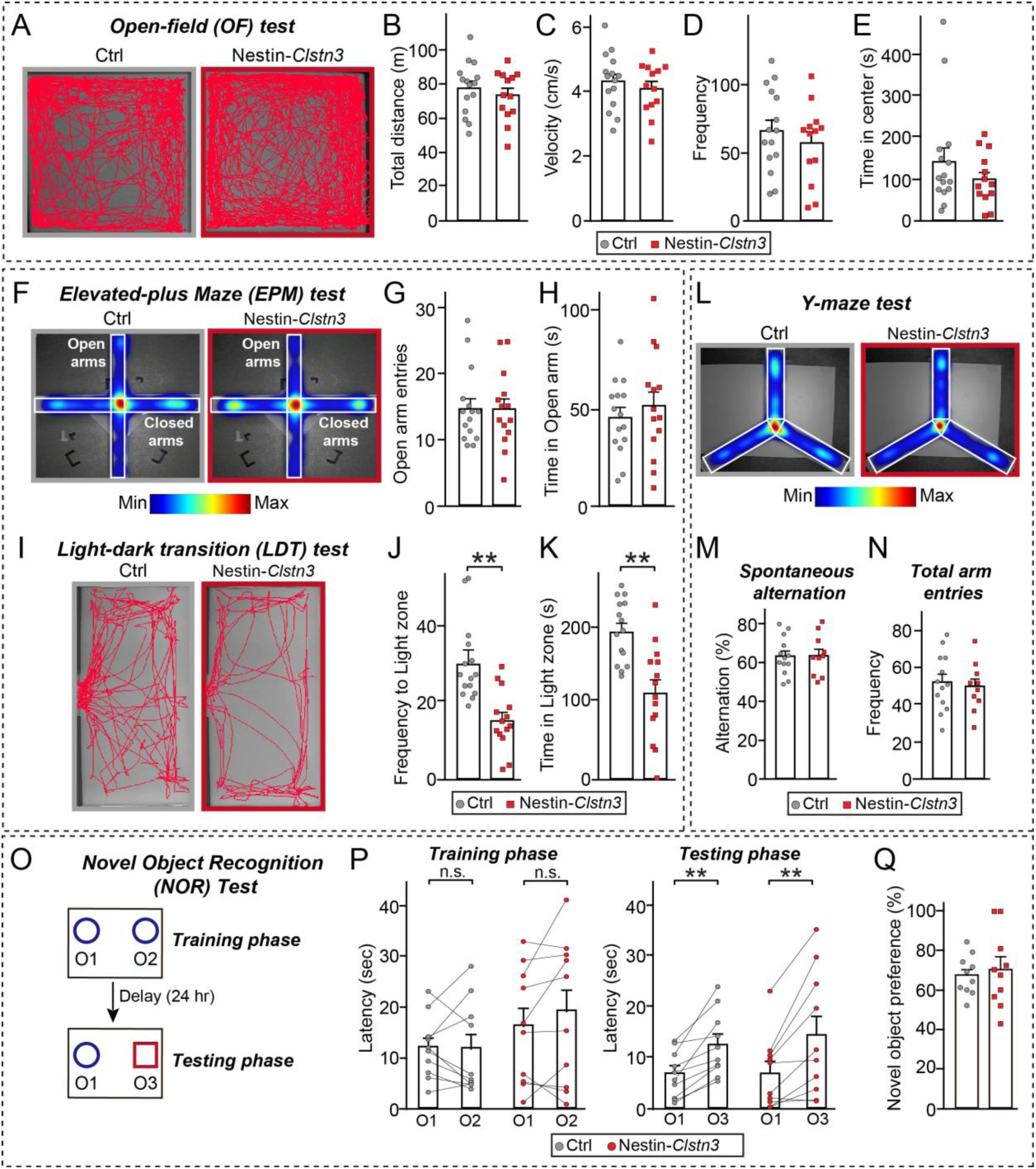
Deletion of Clstn3 increases bright light-induced anxiety-related behavior in mice. **(A–E)** Representative heat map images in the open-field test (OFT; **A**) and summary graphs showing quantification of total distance (**B**), velocity (**C**), frequency of entering the center (**D**), and time spent in the center (**E**) recorded from *Clstn3^fl/fl^* (Ctrl) and Nestin*-Clstn3* mice. Data are shown as means ± SEMs (n = 12 mice for Ctrl and n = 9 mice for Nestin*-Clstn3*; n.s., not significant). **(F–H)** Representative heat map images for the elevated-plus maze (EPM; **F**) and summary graphs showing quantification of the number of open arm entries (**G**) and time in open arms (**H**). Data are shown as means ± SEMs (n = 12 mice for Ctrl and n = 9 mice for Nestin*-Clstn3* mice). **(I–K)** Representative heat map images for the light zone during the light-dark transition test (LDT; **I**) and summary graphs showing quantification of transition frequency (**J**) and time spent in the light zone (**K**) recorded from *Ctrl* and Nestin*-Clstn3*) mice. Data are shown as means ± SEMs (n = 12 mice for Ctrl and n = 9 mice for Nestin*-Clstn3* mice; \*\**p* < 0.01, Mann-Whitney U test). (**L–N**) Normal spatial recognition memory of Nestin*-Clstn3* mice in the Y-maze. Representative heat map images (**L**) and summary graphs showing quantification of spontaneous alternation (**M**) and total arm entries (**N**). Data are shown as means ± SEMs (n = 12 mice for Ctrl and n = 9 mice for Nestin*-Clstn3* mice). **(O**–**Q)** Schematic depiction of the novel object-recognition test (NOR; **O**) and summary graphs showing quantification of preference for a novel object (**P** and **Q**). Data are shown as means ± SEMs (n = 12 mice for Ctrl and n = 9 mice for Nestin*-Clstn3* mice).

### Cadherin domains of Clstn3 are required for restoration of the impaired excitatory innervations observed in Clstn3-cKO hippocampal neurons

To investigate the physiological significance of Clstn3–Nrxn interactions *in vivo*, we constructed AAVs encoding full-length Clstn3 (Full), cadherin domains plus intracellular regions (Cad), or a cadherin domains-deleted Clstn3 fragment (ΔCad) (***Figure 7A*** and ***Figure 7—figure supplement 1***). We then stereotactically injected AAVs expressing the indicated Clstn3 protein into the CA1 hippocampus of ∼8-wk-old *Clstn3^tm1a/tm1a^* or Nestin-*Clstn3* mice and performed quantitative immunofluorescence analyses after 2 wks using antibodies to VGLUT1 or PSD-95 (an excitatory postsynaptic marker) (***Figures 7B–E*, *Figure 5— figure supplement 1*, *Figure 7—figure supplement 1*** and ***2***). Strikingly, expression of Clstn3 Full or Clstn3 Cad, but not Clstn3 ΔCad, restored the decreased VGLUT1 puncta density to WT mouse levels in the SR layers in both *Clstn3^tm1a/tm1a^* or Nestin-*Clstn3* mice (***Figures 7A–E*, *Figure 5—figure supplement 1***). Expression of Clstn3 Full also restored the decreased VGLUT1 puncta density in the SO layer, but expression of neither Clstn3 Cad nor Clstn3 ΔCad exhibited rescuing effect in both *Clstn3^tm1a/tm1a^* or Nestin-*Clstn3* mice (***Figures 7A–E*, *Figure 5—figure supplement 1***). In contrast, the decreased PSD-95 puncta density was rescued by expression of Clstn3 Cad or Clstn3 ΔCad in both SO and SR layers of Nestin-*Clstn3* mice (***Figure 7—figure supplement 2***). Taken together, these results suggest that Clstn3 organizes the development of hippocampal CA1 excitatory synapses, and likely acts through cadherin domains-mediated interactions with presynaptic Nrxns to control the properties of specific excitatory synaptic projections involving the SR layer.

**Figure 7.**
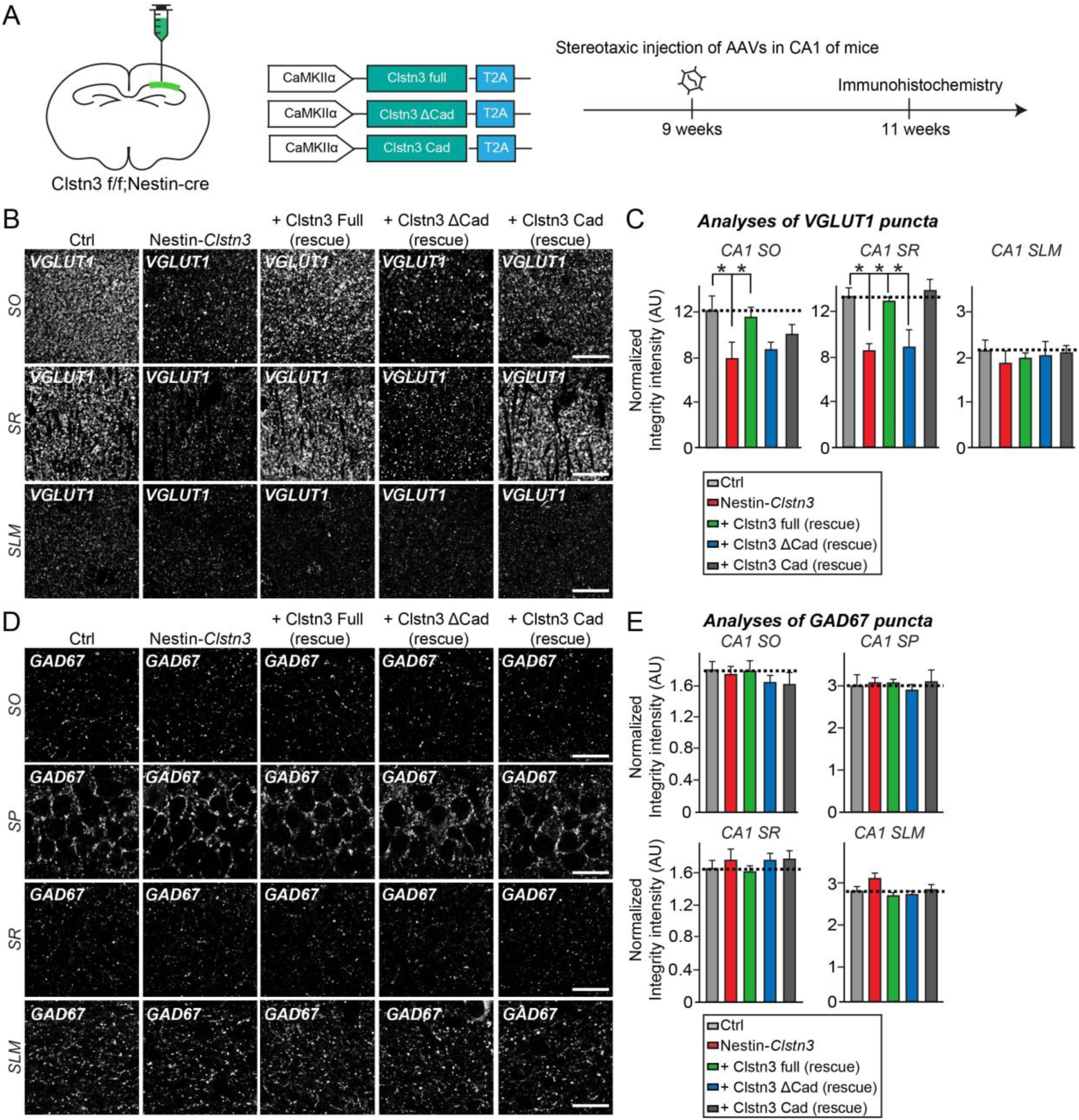
Nrxn Interactions are required for Clstn3-mediated regulation of specific Schaffer-collateral projections in the CA1 hippocampal area. **(A)** Experimental protocols for quantitative immunohistochemistry experiments presented in (**B–E**). Quantitative immunohistochemical analyses were performed 2 wk after stereotactic injection of the indicated AAVs into the CA1 hippocampal regions of ∼9-wk-old Nestin-*Clstn3* mice. **(B)** Representative images of hippocampal CA1 SO, SR and SLM regions 2 wk after stereotactic injections of the indicated AAVs into *Clstn3^fl/fl^* (Ctrl) or Nestin-*Clstn3* mice, followed by immunostaining for the excitatory synapse marker VGLUT1. Scale bar: 20 μm (applies to all images). **(C)** Quantification of the integrated intensity of VGLUT1-positive synaptic puncta. Data are means ± SEMs (\**p* < 0.05; non-parametric Kruskal-Wallis test with Dunn’s *post hoc* test; n = 5 mice each after averaging data from 6 sections/mouse). **(D)** Representative images of hippocampal CA1 SO, SP, SR and SLM regions 2 wk after stereotactic injection of the indicated AAVs into *Clstn3^fl/fl^* or Nestin-*Clstn3* mice, followed by immunostaining for the inhibitory synapse marker GAD67. Scale bar: 20 μm (applies to all images). **(E)** Quantification of the integrated intensity of GAD67-positive synaptic puncta. Data are means ± SEM (non-parametric Kruskal-Wallis test with Dunn’s *post hoc* test; n = 5 mice each after averaging data from 6 sections/mouse).

## Discussion

The present study was initiated with the goal of reconciling discrepancies surrounding molecular mechanisms of Clstn3-mediated synapse development. For this, we revisited the most critical (and controversial) issues using newly engineered Nrxns expression vectors and recombinant proteins, and a newly developed Clstn3-cKO mice. Taken together with our previous findings, the current study’s three principal observations provide plausible explanations for discrepancies surrounding the molecular mechanisms of Clstn3-mediated synapse development.

First, we found that Clstn3 binds directly to β-Nrxns (***Figures 1*** an*d **2***). This interaction was only detected using newly engineered Nrxn1β constructs that were expressed at ∼100-fold higher levels (***Figures 1, 2,*** and ***Figure 1—figure supplement 1***). Because Nrxn1β dStalk1 displayed significant decrease binding to Clstn3 and this juxtamembrane sequence is heavily glycosylated, we initially speculated that glycosylation patterns contributed to Clstn3 binding (***Figure 2*** and ***Figure 2—figure supplement 1***); however, removal of glycosylated residues did not impair binding (***Figure 2—figure supplement 1***). Instead, we conclude that this sequence may influence the orientation and/or conformation of the LNS domain in a manner that is critical for its interaction with Clstn3. Structural studies have suggested that a β-Nrxn LNS domain containing an SS4 insertion exists as a dynamic equilibrium between two conformational states: one in which the splice insert forms a protruded α-helix that supports binding to cerebellins (Cblns), and one in which the additional residues adopt a β-sheet conformation and restore binding to NLs and LRRTMs (Reissner et al., 2013). Because Clstn3 exhibits a slight preference for binding SS4-positive Nrxns, it is tempting to speculate that restoration of downstream residues promoted a conformational change in the SS4 insert, leading to Clstn3 binding. As noted above, the current study, unlike our previous study and that of Pettem et al., employed newly engineered Nrxn recombinant proteins to provide the first demonstration of direct interactions with recombinant Clsn3 proteins (***Figure 2***). Moreover, in keeping with results from the current study, it was previously reported that β1-Nrxn binds immobilized Clstn3 with a complex binding mode (Lu et al., 2014). Although it was claimed that Clstn3 interacts with α-Nrxns (but not β-Nrxns), and we previously failed to detect the interaction of Nrxns with Clstn3 (Pettem et al., 2013; Um et al., 2014), we would argue that conclusions based on binding experiments employing a limited set of methodologies could unintentionally be misleading, as was the case for our previous report (Um et al., 2014).

Second, re-expressing select Nrxn1 variants in Nrxn-deficient neurons rescued impaired Clstn3 synaptogenic activity (***Figure 3***). Previously, we reported that presynaptic Nrxns serve as functional receptors for postsynaptic Clstn3, but do not directly interact with them (Um et al., 2014). Our new data indicate that our previous model should incorporate direct binding of β-Nrxns to Clstn3 (***Figures 1–3***). In support of this interpretation, BAM-2, a Nrxn-related *C. elegans* homolog, was recently reported to bind CASY-1 and mediate neural circuit wiring of male-specific hook-sensory HOA neurons in *C. elegans* (Kim and Emmons, 2017). Intriguingly, CASY-1 does not directly interact with NRX-1, a canonical *C. elegans* Nrxn ortholog (Kim and Emmons, 2017). Although BAM-2 exhibits considerable sequence homology/similarity with α-Nrxns (Colavita and Tessier-Lavigne, 2003), whether these vertebrate and worm genes are functionally homologous remains to be determined.

Nrxns interact with various postsynaptogenic proteins, including NLs, LRRTMs, Cblns, and latrophilins (Südhof, 2017). These interactions are dynamically modulated by the alternative splicing status of Nrxns, mainly at the canonical SS4 splice site. Strikingly, the synapse-promoting activity of Clstn3 required Nrxn splice variants containing the SS4 insert (***Figure 3***), as has been shown for other Nrxn ligands (i.e., Cblns) (Südhof, 2017). These observations are consistent with slightly stronger binding of Nrxn1β^+SS4^ to Clstn3, which contrasts with the exclusive binding of Nrxn1β^+SS4^ to Cblns (***Figures 1G*** and ***1H***). Pull-down experiments in mouse brains also showed that Clstn3 is primarily associated with Nrxn-SS4–positive variants *in vivo* and that its synaptogenic activity requires the binding to Nrxn-SS4–positive variants (***Figures 3C*** and ***3D***) (Um et al., 2014). Additionally, the shed ectodomain of Clstn3 suppresses the synaptogenic activity of NL-2 and LRRTM2, suggesting that Clstn3 competes with these Nrxn ligands (Pettem et al., 2013). However, LRRTM2 interacts only with Nrxn-SS4–negative splice variants (Ko et al., 2009a), whereas Clstn3 activity requires Nrxn-SS4–positive splice variants; moreover, a β-Nrxn LNS domain (identical to the sixth LNS domains-sixth LNS domain in α-Nrxns) is sufficient for Clstn3 binding. Although we still failed to detect clear interactions of α-Nrxns with Clstn3 in cell surface-binding assays (***data not shown***), we observed robust Nrxn1α–Clstn3 interactions in brain pull-down and direct binding assays (***Figures 2E, 3C*** and ***3D***). It is possible that the surface-binding assays employed here were too stringent to observe this interaction, or the cell surface presentation of α-Nrxns was not optimal for binding. Thus, we conclude that Clstn3 binds to both α- and β-Nrxns through a common LNS domain, but a subset of Clstn3 synaptic functions might require other unidentified ligands *in vivo*. Moreover, given the activity-dependent regulation of Nrxn alternative splicing at the SS4 site (Iijima et al., 2011), future studies should investigate whether Clstn3–Nrxn interactions could be fine-tuned depending on distinct activity patterns *in vivo*. Furthermore, given a recent report that different SS4-positive splice variants of two different Nrxns (Nrxn1 vs. Nrxn3) differentially control different postsynaptic responses (Dai et al., 2019), it is possible that Clsnt3 preferentially partners with a specific Nrxn *in vivo*.

Third, re-expressing Clstn3 cadherin domains was sufficient to completely rescue the impaired excitatory synapse structures in hippocampal CA1 neurons from both Clstn3*^tm1a/tm1a^* and Nestin-*Clstn3* mice (***Figure 7*** and ***Figure 5—figure supplement 1***), but only in the SR layer (but see ***Figure 7—figure supplement 2***). In contrast, the structural deficits in other hippocampal CA1 layers were rescued by expression of full-length Clstn3 protein, but not by expression of partial Clstn3 proteins (***Figure 7*** and ***Figure 5—figure supplement 1***). Although it is possible that other unidentified proteins could also bind to Clstn3 cadherin domains, it is likely that presynaptic neurexins and postsynaptic Clstn3 control the properties of excitatory synapse development in specific Schaffer-collateral projections within the SR layer. A recent paper highlighted the fact that the distinctive features of two different Schaffer-collateral projections differentially regulate mushroom spine density and high-magnitude LTP in the SO layer, organized by heterophilic type II cadherins (Basu et al., 2017). Moreover, Nrxn genes show differential, but overlapping, isoform- and region-dependent expression in different classes of neurons, and undergo highly distinctive, cell type-specific alternative splicing (Nguyen et al., 2016; Ullrich et al., 1995). One plausible scenario would be that CA3 neurons projecting to CA1 neurons in the SR layer, but not the SO layer, express higher levels of SS4-positive Nrxns at their nerve terminals. A recent extensive chromogenic and fluorescent *in situ* hybridization analysis showed that the levels of Nrxn mRNAs in hippocampal subfields are significantly lower in GABAergic neurons than excitatory neurons (Uchigashima et al., 2019), which might account for preferential deficits on excitatory synapses in CA1 hippocampal regions of Clstn3 KO mice. Another possible scenario is that major glutamatergic axon fibers of Schaffer-collateral (SC) pathways are targeted to CA1 pyramidal neurons in the SR layer, whereas a subpopulation of axon fibers of SC pathways innervate both bistratified GABAergic interneurons in the SO layer and CA1 pyramidal neurons. Additional studies will also be required to determine the identity of neurons *in vivo* that are responsible for Nrxn–Clstn3 interactions in other neural circuits inside and outside of the hippocampus. Notably, in stark contrast to the previous report that decreased inhibitory synapse structure and transmission in Clstn3-KO mice (Pettem et al., 2013), we found that inhibitory synapses in CA1 hippocampal layers and the cortex were morphologically normal in our Clstn3-KO mice (***Figure 5*** and ***Figure 5—figure supplement 1***). Moreover, to our surprise, Clstn3-KO mice did not exhibit any alterations in basal synaptic transmission at either excitatory or inhibitory synapses (***Figure 5—figure supplement 2***), suggesting that Clstn3 is required for structural integrity, but not basal synaptic transmission, in hippocampal CA1 neurons. The reason for this discrepancy is currently unclear, but it is likely that Clstn family proteins are functionally redundant in the maintenance of basal synaptic transmission; alternatively, Clstn3 may be specifically required for certain forms of synaptic plasticity. A notable difference between the study of Pettem et al. and the current study is that the former targeted exons 2 and 3 of the mouse *Clstn3* gene, whereas our study targeted exon 8 (***Figure 4A***). In addition, Pettem *et al* employed an EIIa-Cre driver line that permits germ line deletion in both excitatory and inhibitory neurons by driving expression of Cre recombinase in the early mouse embryo, whereas we used a pan-neuronal Nestin-Cre driver line that is presumed to be similar to EIIa-Cre. Because Clstn3 is expressed in both excitatory and inhibitory neurons (***Figure 4—figure supplement 2***), it is possible that specifically deleting Clstn3 in GABAergic inhibitory neurons may produce marked deficits in GABAergic synapse development, as previously documented (Pettem et al., 2013), but this did not clearly manifest in our Clstn3-cKO mice. A more systematic, follow-up investigation to probe cell type-specific contributions of Clstn3 to distinct anatomical and electrophysiological phenotypes should address these important, but puzzling, observations. Future studies should also probe how Nrxn–Clstn3 complexes mediate synaptic specificity involving the SR layer of the hippocampal CA1 region. Furthermore, rigorous analyses are warranted to address whether and how this circuit specificity is related to the altered anxiety-like behavior observed in Clstn3-cKO mice (***Figure 6***). A complete understanding of how Clstn3 functions will also require investigation of the potential contributions of other Clstn3 domains (e.g., LNS domain, intracellular region) to the functions of Clstn3, as was similarly shown in *C. elegans* (Ikeda et al., 2008; Kim and Emmons, 2017; Thapliyal et al., 2018).

In summary, the present study unambiguously establishes that Clstn3 plays a role in specifying the properties of a specific hippocampal CA1 neural circuit, in part by regulating excitatory synapse development through formation of physical complexes with specific Nrxn splice variants.

## Materials and methods

### Plasmids

<u>Nrxn rescue vectors</u> were generated by PCR-amplification of full-length sequences of mouse Nrxn1β^-SS4^, Nrxn1β^+SS4^, rat Nrxn1β^-SS4^, rat Nrxn1β^+SS4^, bovine Nrxn1α^-SS4^ and bovine Nrxn1α^+SS4^, followed by digestion with *Nhe*I and *BsrG*I and cloning into the TKD vector [L-313 vector]. Three nucleotides (underlined) in the GTGCC<u>T</u>TC<u>C</u>TC<u>T</u>ATGACAACT sequence were then mutated to render it shRNA-resistant. pGW1-FLAG-mNrxn1β^-SS4^ and pGW1-FLAG-mNrxn1β^+SS4^ were generated by PCR-amplification of full-length mNrxn1β^-SS4^ and mNrxn1β^+SS4^, respectively, digestion with *Kpn*I and *EcoR*I, and cloning into a modified pGW1 vector containing a mouse Clstn1 signal peptide and FLAG epitope (pGW1 vector; British Biotechnology, Oxford, UK). <u>pCMV-IgC-</u>mNrxn1β^-SS4^ and pCMV-IgC-mNrxn1β^+SS4^ were generated by PCR-amplification of the indicated extracellular regions of mNrxn1β^-SS4^ (aa 1–359) and mNrxn1β^+SS4^ (aa 1–389), respectively, followed by digestion with *EcoR*I and *Sal*I and cloning into a pCMV-IgC vector. <u>pGW1-FLAG-Clstn3</u> and <u>pGW1-FLAG-Clstn3-ΔCAD</u> were generated by PCR-amplification of full length mouse Clstn3 and a Clstn3 fragment (aa 258–956), respectively, followed by digestion with *Kpn*I and *EcoR*I and cloning into the pGW1-FLAG vector. <u>pGW1-FLAG-Clstn3-Cad</u> used in **Figure 2A** was generated by PCR amplification of two Clstn3 fragments (aa 29–252 and aa 848–956), followed by digestion with *Kpn*I and *EcoR*I for the first fragment and *EcoR*I only for the second fragment, and subsequent cloning into the pGW1-FLAG vector. The following deletion constructs of mNrxn1β were generated using pGW1-FLAG-mNrxn1β^+SS4^: <u>pGW1-FLAG-mNrxn1β-ΔLNS6</u> was generated by PCR amplification of mNrxn1β (aa 293–468), followed by digestion with *Kpn*I and *EcoR*I and cloning into the pGW1-FLAG vector. <u>pGW1-FLAG-mNrxn1β-ΔStalk1</u> was generated by PCR amplification of two mNrxn1β fragments (aa 1–315 and aa 347–468), followed by digestion with *Kpn*I and *EcoR*I for the first fragment and *EcoR*I for the second fragment, and subsequent cloning into the pGW1-FLAG vector. <u>pGW1-FLAG-mNrxn1β-ΔStalk2</u> was generated by PCR amplification of two mNrxn1β fragments (aa 1–298 and aa 388-468), followed by digestion with *Kpn*I and *EcoR*I for the first fragment and *EcoR*I only for the second fragment, and subsequent cloning into the pGW1-FLAG vector. <u>pGW1-FLAG-mNrxn1β-ΔCysL</u> and <u>pGW1-FLAG-mNrxn1β-ΔCHO</u> were modified from the previously described mNrxn1β point mutants [see (Sterky et al., 2017) and **Figure 2**] by replacing aa 299–468 of WT mNrxn1β with the identical region from the mutated mNrxn1β and cloning the resulting sequence into the pGW1-FLAG vector using *Sal*I and *EcoR*I sites. <u>pGW1-FLAG-mNrxn1β-ΔHRD</u> was generated by PCR amplification of mNrxn1β (aa 84-468), followed by digestion with *Kpn*I and *EcoR*I and cloning into the pGW1-FLAG vector. pCAGG-mNrxn1 β^+SS4^-FLAG-ΔHS, L-313-mNrxn1β^-SS4^-ΔHS-<u>FLAG</u> and L-313-mNrxn1β^+SS4^-ΔHS-FLAG were generated by site-directed mutagenesis to change a single amino acid from ILVA<u>S</u>AECP to ILVA<u>A</u>AECP (underlined residues denote the changed residue) using pCAGG-mNrxn1 β^+SS4^-FLAG, L-313-mNrx1β^-SS4^-FLAG, and L-313-mNrxn1β^+SS4^-FLAG as templates, respectively. <u>pGW1-FLAG-mNrxn1γ</u> was generated by PCR amplification of full-length mNrxn1γ, followed by digestion with *Kpn*I and *EcoR*I and cloning into the pGW1-FLAG vector. Constructs of the extracellular domain of mouse Clstn3 (Clstn3 ecto: aa 21–846; Clstn3 Cad: aa 21–269; Clstn3 ΔCAD: aa 258–846) used in **Figure 2D** were N-terminally fused to a hexa-histidine tag and HA tag and cloned into the pCAGGS vector using *EcoR*I and *Not*I restriction enzymes. The following constructs were previously described: <u>L-315-Nrxn-TKD</u> (Um et al., 2016); <u>pCMV5-mVenus-NL-1</u> (Lee et al., 2013); <u>pCMV-IgC-Clstn3</u> (Um et al., 2014b); <u>pCMV-IgC-NL-2</u> (Lee et al., 2013); <u>pDisplay-LRRTM2 (Ko et al., 2011)</u>; pCAGG-mNrxn1α^-SS4^-FLAG, <u>pCAGG-</u>mNrxn1α^+SS4^-FLAG, pCAGG-mNrxn1β^-SS4^-FLAG, pCAGG-mNrxn1β^+SS4^-FLAG, <u>pCMV-IgC-</u>mNrxn1α^-SS4^ and pCMV-IgC-mNrxn1α^+SS4^ (Matsuda and Yuzaki, 2011). <u>pAAV2-CAG-Clstn3 Full</u>, <u>pAAV2-CAG-Clstn3 Cad</u> and <u>pAAV2-CAG-Clstn3 ΔCad</u> were generated by PCR-amplification using pGW1-FLAG-Clstn3 Full, pGW1-FLAG-Clstn3 Cad, and pGW1-Flag-Clstn3 ΔCad as templates, respectively, followed by digestion with *EcoR*V and *Hind*III and cloning into a pAAV_2_-CAG vector.

### Antibodies

Fusion proteins of glutathione-S-transferase and mouse Clstn3 (JK091; aa 21–244) were produced in BL21 *Escherichia coli* and purified using a glutathione-Sepharose column (GE Healthcare, Chicago, IL, USA). Clstn1 peptides (JK025; CHQRTMRDQDTGKEN) and Clstn2 peptides (JK028; CSSQSPERSTWNTAGVINIWK) were synthesized and conjugated to keyhole limpet hemocyanin through a cysteine added to the N-terminus of the peptide. Following immunization of rabbits with the immunogen, antibodies were affinity-purified using a Sulfolink column (Pierce). The following commercially available antibodies were used: mouse monoclonal anti-HA (clone HA-7; Covance), mouse monoclonal anti-FLAG M2 (Sigma-Aldrich, St. Louis, MO, USA), rabbit polyclonal anti-FLAG (Sigma-Aldrich), goat polyclonal anti-EGFP (Rockland Immunochemicals), mouse monoclonal anti-NL1 (clone N97A/31; NeuroMab), rabbit polyclonal anti-NL2 (Synaptic Systems, Göttingen, Germany), rabbit monoclonal anti-TrkC (clone C44H5; Cell Signaling Technology), guinea pig polyclonal anti-VGLUT1 (Millipore), mouse monoclonal anti-GAD67 (clone 1G10.2; Millipore), mouse monoclonal anti-PSD-95 (clone K28/43; Thermo-Fisher), mouse monoclonal anti-β-actin (clone AC-74; Sigma-Aldrich), mouse monoclonal anti-parvalbumin (clone PARV-19; Millipore), mouse monoclonal anti-gephyrin (clone 3B11; Synaptic Systems), rabbit polyclonal anti-GABAA receptor γ2 (Synaptic Systems); mouse monoclonal anti-GluN1 (clone 54.1; Millipore), rabbit polyclonal anti-GluN2A (Millipore), rabbit polyclonal anti-GluN2B (Millipore), goat anti-human IgG-peroxidase (Sigma-Aldrich), and mouse monoclonal anti-NeuN (clone A60; Millipore). The following antibodies have been previously described: anti-PSD-95 [JK016] (Um et al., 2016), anti-synapsin [JK014] (Han et al., 2016), anti-Clstn3 [JK001] (Um et al., 2014b), anti-GluA1 [1193], and anti-GluA2 [1195] (Kim et al., 2009).

### Animals

*Clstn3^tm1a(EUCOMM)Hmgu^* mice, in which a targeting cassette harboring FRT, lacZ and loxP sites is inserted between exon 7 and exon 8, resulting in a ‘knockout-first’ lacZ reporter-tagged *Clstn3^tm1a^* insertion allele with conditional potential, were obtained from the European Mouse Mutagenesis Consortium (EUCOMM). To generate conditional *Clstn3* KO mice (*Clsnt3^flox/flox^*), we first bred *Clsnt3^tm1a/tm1a^* mice in a C57BL/6J background with FLPO recombinase driver mice [C57BL/6N-Tg(CAG-Flpo)1Afst/Mmucd, MMRRC] to generate *Clsnt3^tmc1c/tm1c^* mice. Male homozygous *Clsnt3^tmc1c/tm1c^* mice in a C57BL/6J genetic background were crossed with female heterozygous *Clsnt3^tm1c/+^* mice (for Nestin conditional) carrying the Cre transgene. A Nestin-Cre (003771) line in a C57BL/6N genetic background, generated by crossing the original Cre-driver lines purchased from the Jackson Laboratory with C57BL/6J for more than five generations, was obtained from Dr. Eunjoon Kim (KAIST, Korea). Cre-negative *Clsnt3^flox/flox^* littermates were used as controls in all experiments. Mice were housed and bred at the animal facility of Daegu Gyeongbuk Institute of Science and Technology (DGIST), and their experimental uses were approved by the Institutional Animal Care and Use Committee of DGIST (DGIST-IACUC-19052110-00). Mice were given *ad libitum* access to food and water, and were maintained in a controlled environment with a 12-h light/dark cycle. Mice were weaned at the age of P24–27, and mixed-genotype littermate mice were group-housed (4–5 mice per cage) until experiments. Genotyping was performed on genomic DNA extracted from tail biopsies using the 2X Taq PCR Smart mix 2 kit (Solgent Co.). Genotyping primers were as follows: Clstn3 forward, 5’-ACT TGA TCA GTC CTC CTG CAT CAG-3’; Clstn3 reverse, 5’-CTG AAG TTC AGG GTC AGC CTG TAA-3’; FRT reverse, 5’-CCT TCC TCC TAC ATA GTT GGC AGT-3’; Cre [Nestin-Cre] forward, 5’-CCG CTT CCG CTG GGT CAC TGT-3’; WT reverse, 5’-CTG AGC AGC TGG TTC TGC TCC T-3’; and Cre [Nestin-Cre] reverse, 5’-GAC CGG CAA ACG GAC AGA AGC A-3’. A PCR product of ∼130 base pairs (bp) was obtained from WT DNA, whereas *Clsnt3^flox^* produced a PCR product of ∼300 bp. The Cre transgene from the Nestin-cre mouse produced a PCR product of ∼ 400 bp.

### Production of recombinant lentiviruses and adeno-associated viruses (rAAVs)

#### 1. Lentivirus production

Recombinant lentiviruses were produced as previously described (Lee et al., 2013). In brief, HEK293T cells were transfected with three plasmids—Lentivirus vectors, psPAX2, and pMD2G—at a 2:2:1 ratio using Lipofectamine 2000 (Thermo-Fisher Scientific) according to the manufacturer’s protocol. After 72 h, lentiviruses were harvested by collecting the medium from transfected HEK293T cells and briefly centrifuging at 1,000 × g to remove cellular debris. Filtered media containing 5% sucrose were centrifuged at 117,969 × g for 2 h; supernatants were then removed and the virus pellet was washed with ice-cold phosphate-buffered saline (PBS) and resuspended in 80 μl PBS.

#### 2. AAV production

HEK293T cells were co-transfected with the indicated AAV vectors and pHelper and pRC1-DJ vectors. Seventy-two hours later, transfected HEK293T cells were collected, lysed, and mixed with 40% polyethylene glycol and 2.5 M NaCl, and centrifuged at 2000 × g for 30 min. The cell pellets were resuspended in HEPES buffer (20 mM HEPES; 115 mM NaCl, 1.2 mM CaCl_2_, 1.2 mM MgCl_2_, 2.4 mM KH_2_PO_4_) and an equal volume of chloroform was added. The mixture was centrifuged at 400 × g for 5 min, and concentrated three times with a Centriprep centrifugal filter (Millipore) at 1,220 × g for 5 min each and with an Amicon Ultra centrifugal filter (Millipore) at 16,000 × g for 10 min. Before titering AAVs, contaminating plasmid DNA was eliminated by treating 1 μl of concentrated, sterile-filtered AAVs with 1 μl of DNase I (Sigma-Aldrich) for 30 min at 37°C.

After treatment with 1 μl of stop solution (50 mM ethylenediaminetetraacetic acid) for 10 min at 65°C, 10 μg of protease K (Sigma-Aldrich) was added and AAVs were incubated for 1 h at 50°C. Reactions were inactivated by incubating samples for 20 min at 95°C. The final virus titer was quantified by qRT-PCR detection of *EGFP* sequences and subsequent reference to a standard curve generated using the pAAV-U6-EGFP plasmid. All plasmids were purified using a Plasmid Maxi Kit (Qiagen GmbH).

### LC-MS/MS protein analysis

Peptides were analyzed using a nanoflow LC-MS/MS system consisting of an Easy nLC 1000 system (Thermo-Fisher) and an LTQ Orbitrap Elite mass spectrometer (Thermo-Fisher) equipped with a nano-electrospray source. Peptide solutions (5-µl aliquots) were loaded onto a C_18_ trap column (20 × 75 µm, 3 µm particle size; Thermo-Fisher) using an autosampler. Peptides were desalted and concentrated on the column at a flow rate of 5 µl/min. Trapped peptides were then separated on a 150-mm custom-built microcapillary column consisting of C_18_ (particle size, 3 µm; Aqua Science, Yokohama, Japan) packed in 100-µm silica tubing with a 6-µm inner diameter orifice. The mobile phases A and B were composed of 0% and 100% acetonitrile, respectively, and each contained 0.1% formic acid. The LC gradient began with 5% B for 5 min and was increased to 15% B over 5 min, 50% B over 55 min, and 95% B over 5 min, then remained at 95% B over 5 min, followed by 5% B for an additional 5 min. The column was re-equilibrated with 5% B for 15 min between runs. A voltage of 2.2 kV was applied to produce the electrospray. In each mass analysis duty cycle, one high-mass resolution (60,000) MS spectrum was acquired using the orbitrap analyzer, followed by 10 data-dependent MS/MS scans using the linear ion trap analyzer. For MS/MS analysis, a normalized collision energy (35%) was used throughout the collision-induced dissociation phase. All MS and MS/MS spectra were acquired using the following parameters: no sheath and auxiliary gas flow; ion-transfer tube temperature, 200°C; activation Q, 0.25; and activation time, 20 ms. Dynamic exclusion was employed with a repeat count of 1, a repeat duration of 30 s, an exclusion list size of 500, an exclusion duration of 60 s, and an exclusion mass width of ± 1.5 m/z.

### MS data analysis

MS/MS spectra were analyzed using the following analysis protocol, referencing the UniProt mouse database (09-15-2015 release). Briefly, each protein’s reversed sequence was appended onto the database to calculate the false discovery rate. Peptides were identified using ProLuCID (Xu et al., 2015) in Integrated Proteomics Pipeline software, IP2 (http://www.integratedproteomics.com), with a precursor mass error of 25 ppm and a fragment ion mass error of 600 ppm. Trypsin was used as the protease, and two potential missed cleavages were allowed. Carbamidomethylation at cysteine was chosen as a static modification, and methionine oxidation was chosen as a variable modification. Protein lists consisting of two or more peptide assignments for protein identification (false-positive rate < 0.01) were prepared by filtering and sorting output data files.

### Primary neuron culture, immunocytochemistry, image acquisition, and quantitative analyses

The indicated analyses were performed using cultured, E18-derived, rat hippocampal neurons and confocal microscopy, as previously described (Kang et al., 2016; Um et al., 2014a; Um et al., 2014b). Rat hippocampal neurons were prepared from E18 rat brains and cultured on coverslips coated with poly-D-lysine in Neurobasal media supplemented with B-27 (Thermo-Fisher), 0.5% fetal bovine serum, 0.5 mM GlutaMax (Thermo-Fisher), and sodium pyruvate (Thermo-Fisher). For immunocytochemistry, cultured neurons were fixed with 4% formaldehyde/4% sucrose, permeabilized with 0.2% Triton X-100 in PBS, and immunostained with primary antibodies and Cy3-or fluorescein isothiocyanate (FITC)-conjugated secondary antibodies (Jackson ImmunoResearch, West Grove, PA, USA). Images were acquired using a confocal microscope (LSM700; Carl Zeiss) equipped with a 63× objective lens. All image settings were kept constant. Z-stack images were converted to maximal projection. All images were quantitatively analyzed in a blinded manner using MetaMorph software (Molecular Devices).

### Cell-surface–binding assays

Ig-fusion proteins of Clstn3, Ig-Nrxn1β splice variants, and IgC alone (Control) were produced in HEK293T cells. Soluble Ig-fused proteins were purified using protein A-Sepharose beads (GE Healthcare). Bound proteins were eluted with 0.1 M glycine (pH 2.5) and immediately neutralized with 1 M Tris-HCl (pH 8.0). Transfected HEK293T cells expressing the indicated plasmids were incubated with 10 μg/ml Ig-fused proteins for 2 h at 37°C. Images were acquired using a confocal microscope (LSM700; Carl Zeiss).

### Pull-down assays

HEK293T cells were transfected with pGW1-FLAG-Clstn3 Full, Cad, ΔCad or pCMV-mVenus-NL-1, harvested 48 h later, and incubated for 1 h at 4°C in solubilization buffer (25 mM Tris-HCl pH 7.6, 150 mM NaCl, 1% NP-40, 1% sodium deoxycholate, 0.1% sodium dodecyl sulfate, 5 mM CaCl_2_, 5 mM MgCl_2_). The suspension was centrifuged at 20,000 × g to remove insoluble debris, and each supernatant was mixed with 10 μg of IgC (control), Ig-Nrxn1α^+SS4^ or Ig-Nrxn1β^+SS4^ supplemented with protein A-Sepharose beads and incubated at 4°C for 2 h with gentle agitation. For *in vivo* pull-down assays, synaptosomal fractions prepared from mouse brains were mixed with 10 μg of IgC (control), Ig-Nrxn1α^-SS4^, Ig-Nrxn1α^+SS4^, Ig-Nrxn1 β^-SS4^ or Ig-Nrxn1β^+SS4^ and incubated at 4°C for 2 h with gentle agitation. Protein A-Sepharose beads were washed three times with solubilization buffer, solubilized in SDS sample buffer, and loaded onto SDS-PAGE gels for immunoblot analyses. The antibodies used for immunoblotting were anti-FLAG (1 μg/ml), anti-EGFP (1:1000), anti-Clstn3 (JK091; 1 μg/ml), anti-NL-1 (1 μg/ml), anti-NL-2 (1 μg/ml), and anti-TrkC (1 μg/ml).

### Direct protein-interaction assays

For direct interaction assays, 10 μg of IgC (control), Ig-Nrxn1α^-SS4^, Ig-Nrxn1α^+SS4^, Ig-Nrxn1 β^-SS4^, or Ig-Nrxn1β^+SS4^ was incubated with 4 μg of purified His-HA-Clstn3 Full, His-HA-Clstn3 Cad, or His-HA-Clstn3 ΔCad for 2 h at 4°C in binding buffer (25 mM Tris pH 7.5, 30 mM MgCl_2_, 40 mM NaCl, and 0.5% Trion X-100). Talon metal affinity resins (Clontech) beads were then added to purified protein mixtures as indicated, and incubated for 2 h at 4°C. Beads were washed three times with binding buffer, solubilized in SDS sample buffer, and loaded onto SDS-PAGE gels for immunoblot analyses. Anti-human IgG was used for immunoblotting.

### Heterologous synapse-formation assays

Heterologous synapse-formation assays were performed using recombinant Clstn3 fusion proteins as previously described (Ko et al., 2009). Briefly, HEK293T cells were transfected with EGFP (negative control) or the indicated Clstn3 constructs using Lipofectamine 2000 (Thermo-Fisher). After 48 h, the transfected HEK293T cells were trypsinized, seeded onto *in vitro* day10 (DIV10) hippocampal neurons, co-cultured for an additional 72 h, and double-immunostained on DIV13 with antibodies against EGFP, HA, and the indicated synaptic markers (synapsin, VGLUT1, or GAD67). All images were acquired using a confocal microscope (LSM700; Zeiss). For quantification, the contours of transfected HEK293T cells were chosen as the region of interest (ROI). Fluorescence intensities of synaptic markers in each ROIs were quantified for both red and green channels using MetaMorph software (Molecular Devices). Normalized synapse density on transfected HEK293T cells was expressed as the ratio of red to green fluorescence.

### Semi-quantitative immunoblot analysis

For semiquantitative immunoblot analyses, brains from P42 WT, *Clstn3^tm1a/ tm1a^*, *Clstn3^fl/fl^*, or *Clstn3^fl/fl^*::Nestin-Cre (Nestin-*Clstn3*) mice were homogenized in 0.32 M sucrose/1 mM MgCl_2_ containing a protease inhibitor cocktail (Thermo-Fisher Scientific) using a Precellys Evolution tissue homogenizer (Bertin Co.). After centrifuging homogenates at 1,000 × g for 10 min, the supernatant was transferred to a fresh microcentrifuge tube and centrifuged at 15,000 × g for 30 min. The resulting synaptosome-enriched pellet (P2) was resuspended in lysis buffer and centrifuged at 20,800 × g, after which the supernatant was analyzed by Western blotting. Quantitation was performed using ImageJ software (National Institutes of Health).

### Immunohistochemistry

WT and *Clstn3^tm1a/tm1a^* mice were transcardially perfused first with PBS and then with 4% paraformaldehyde. After post-fixation overnight, mouse brains were slowly sectioned at 40 μm using a vibratome (VT1200S; Leica) and washed with PBS. Brain sections were incubated in blocking solution containing 10% horse serum, 0.2% bovine serum albumin, and 2% Triton X-100 for 1 h at room temperature (RT), and then incubated overnight at 4°C with primary antibodies against VGLUT1 (1:200), GAD67 (1:100), or NeuN (1:500). After washing three times, sections were incubated with Cy3- or FITC-conjugated secondary antibodies (Jackson ImmunoResearch, West Grove, PA, USA) for 2 h at RT. Sections were then washed extensively, and mounted on glass slides with Vectashield Mounting Medium (Vector Laboratories). Images were acquired by slide scanner microscopy (AxioScan.Z1; Zeiss).

### Nissl staining

WT and *Clstn3^tm1a/tm1a^* mice were perfused first with PBS and then with 4% paraformaldehyde by cardiac injection. Fixed brain tissue was isolated, post-fixed for 12 hours at 4°C, and dehydrated in 30% sucrose for 6–8 days. Thereafter, brain tissue was embedded in OCT (Optimal Cutting Temperature) compound and stored at -80°C. The frozen tissue was mounted and cut at a thickness of 20 μm using a cryostat (Leica CM5120). Slices were mounted on glass slides, washed three times with PBS (15 min each), and permeabilized with 0.1% Triton X-100 in PBS for 10 min.

Permeabilized slices were washed twice with PBS (5 min each), then incubated for 20 min in 200 μl of NeuroTrace 500/525 Green Fluorescent Nissl Stain (Molecular Probes), diluted 1:100 in PBS before use. Thereafter slices were washed in PBS containing 0.1% Triton X-100 for 10 min, then washed twice in PBS (5 min each) and incubated in PBS for 2 h. Slides were dried and mounted using Vectashield Mounting Medium containing 4’,6-diamidino-2-phenylindole (DAPI; Vector Laboratories). Green fluorescence was imaged by slide scanner microscopy (AxioScan.Z1; Zeiss).

### Fluorescent in situ hybridization (RNAscope assay)

Frozen sections (14 µm thick) were cut coronally through the hippocampal formation and thaw-mounted onto Superfrost Plus microscope slides (Advanced Cell Diagnostics). Sections were fixed in 4% formaldehyde for 10 min, dehydrated in increasing concentrations of ethanol for 5 min, air-dried, and then pretreated with protease for 10 min at RT. For RNA detection, sections were incubated in different amplifier solutions in a HybEZ hybridization oven (Advanced Cell Diagnostics) at 40°C. Three synthetic oligonucleotides complementary to the sequence corresponding to nucleotide residues 302-1210 of Mm–Clstn3–tv1, 896-1986 of Mm-CaMKIIα–C2, 18-407 of Mm–SST–C3, and 2-885 of Mm–Pvalb–C3 (Advanced Cell Diagnostics) were used as probes. The labeled probes were conjugated to Alexa Fluor 488, Altto 550 or Altto 647, after which labeled probe mixtures were hybridized by incubating with slide-mounted sections for 2 h at 40°C. Nonspecifically hybridized probes were removed by washing the sections three times for 2 min each with 1X wash buffer at RT, followed by incubation with Amplifier 1-FL for 30 min, Amplifier 2–FL for 15 min, Amplifier 3–FL for 30 min, and Amplifier 4 Alt B–FL for 15 min at 40°C. Each amplifier was removed by washing with 1X wash buffer for 2 min at RT. Slides were imaged with an LSM700 microscope (Zeiss) and analyzed using MetaMorph software (Molecular Devices).

### Stereotaxic injection of rAAVs

Adult (∼8-wk-old) male *Clstn3*^−/−^ or WT littermate mice (C57BL/6 strain) were anesthetized with Avertin (400 mg/kg body weight) by intraperitoneal injection. rAAV solutions (titers ≥ 1 × 10^11^ viral genomes/ml) were injected with a NanoFil syringe (World Precision Instruments) at a flow rate of 0.1 μl/min. The coordinates used for the CA1 region of the dorsal hippocampus were AP -2.5 mm, ML ± 1.5 mm, DV +1.5 mm (from the dura). The site at DV +1.5 mm received a 1-μl injection. Injected mice were allowed to recover for at least 14 d following surgery prior to use in experiments.

### Electrophysiology

Electrophysiological recordings were performed in acute hippocampal CA1 slices. In brief, transverse hippocampal slices were prepared from *Clstn3^tm1d/tm1d^* or WT mice at P35–42, which is 2–3 weeks after stereotactic injection of AAVs encoding Cre, ΔCre, or Clstn3 proteins (Clstn3 Full, Clstn3 Cad, or Clstn3 ΔCad). Mice were anesthetized with isoflurane and decapitated. Their brains were rapidly removed and placed in ice-cold, oxygenated (95% O_2_ and 5% CO_2_) low-Ca^2+/^high-Mg^2+^ solution containing the following: 3.3 mM KCl, 1.3 mM NaH_2_PO_4_, 26 mM NaHCO_3_, 11 mM D-glucose, 0.5 mM CaCl_2_, 10 mM MgCl_2_, and 211 mM sucrose. Slices were equilibrated at 30°C for at least 60 min in oxygenated artificial cerebrospinal fluid (aCSF), consisting of the following: 124 mM NaCl, 3.3 mM KCl, 1.3 mM NaH_2_PO_4_, 26 mM NaHCO_3_, 11 mM D-glucose, 3.125 mM CaCl_2_, and 2.25 mM MgCl_2_. Slices were then transferred to a recording chamber, where they were maintained at 24–27°C and perfused continuously with 95% O_2_- and 5% CO_2_-bubbled aCSF. Whole-cell recordings of miniature postsynaptic currents were performed on CA1 pyramidal neurons voltage clamped at -70 mV. Glass pipettes were filled with an internal solution containing the following: 130 mM CeMeSO_4_, 0.5 mM EGTA, 5 mM TEA-Cl, 8 mM NaCl, 10 mM HEPES, 1 mM QX-314, 4 mM Mg-ATP, 0.4 mM Na-GTP, and 10 mM phosphocreatine-Na2 for mEPSCs; 130 mM CsCl, 1.1 mM EGTA, 2 mM MgCl_2_, 0.1 mM CaCl_2_, 10 mM NaCl, 10 mM HEPES, and 2 mM Na-ATP for mIPSCs. The osmolarity of the internal solution was 290–300 mOsm (pH 7.3; adjusted with CsOH). For mEPSC recordings, 1 μM TTX, 50 μM DL-APV and 100 μM picrotoxin were added to block Na^+^ currents and GABA_A_ receptors, respectively. For miniature inhibitory postsynaptic current (mIPSC) recordings, 1 μM TTX, 20 μM CNQX, 50 μM DL-APV were added to block Na^+^ currents, AMPA and NMDA receptors, respectively. Evoked synaptic responses in the hippocampal CA1 were induced by placing a concentric bipolar electrode at the CA3 striatum radiatum.

### Mouse behaviors

Behavioral experiments were performed in the following order: open-field test, Y-maze test, novel object-recognition test, light/dark transition test, and elevated plus-maze test. All behavioral analyses were performed using male mice. 1. *Y-maze test*. A Y-shaped white acrylic maze with three 40-cm–long arms at a 120° angle from each other was used. Mice were introduced into the center of the maze and allowed to explore freely for 8 min. An entry was counted when all four limbs of a mouse were within the arm. The movement of mice was recorded by a top-view infrared camera, and analyzed using EthoVision XT 10 software. <u>2. *Open field test*.</u> Mice were placed into a white acrylic open-field box (40 × 40 × 40 cm), and allowed to freely explore the environment for 30 min under low-light conditions (∼30 lux). The traveled distance moved and time spent in the center zone by freely moving mice were recorded by a top-view infrared camera, and analyzed using EthoVision XT 10 software (Noldus). <u>3*. Novel object-recognition test*.</u> An open-field maze was used in this test. Mice were habituated to the maze for 10 min. For training sessions, two identical objects were placed in the center of the maze at regular intervals, and mice were allowed to explore the objects for 5 min. After the training session, mice were returned to their home cage for 24 h. For novel object-recognition tests, one of the two objects was exchanged for a new object, placed in the same position of the maze. Mice were returned to the maze and allowed to explore freely for 5 min. The movement of mice was recorded by infrared camera, and the number and duration of contacts were analyzed using EthoVision XT 10 (Noldus). <u>4. *Elevated plus-maze test*.</u> The elevated plus-maze is a plus-shaped (+) white acrylic maze with two open arms (30 × 5 × 0.5 cm) and two closed arms (30 × 5 × 30 cm) positioned at a height of 75 cm from the floor. Light conditions around open and closed arms were ∼300 and ∼30 lux, respectively. For the test, mice were introduced into the center zone of the elevated plus-maze and allowed to move freely for 10 min. All behaviors were recorded by a top-view infrared camera, and the time spent in each arm and the number of arm entries were measured and analyzed using EthoVision XT 10 software (Noldus). <u>5. *Light/Dark transition test*</u> The light/dark box consists of an open-roof, white (light) chamber conjoined to a closed black (dark) chamber with a small entrance to allow free movement between the two chambers. The light chamber was illuminated at 350 lux. The time spent in each chamber and the number of entries were measured and analyzed using EthoVision XT 10 software (Noldus).

### Statistics

All data are expressed as means ± standard errors of the mean (SEM), and significance is indicated with asterisk (compared with a value from control group) or hashtag (compared with a value from experimental group). All experiments were performed using at least three independent cultures and the normality of data distributions was evaluated using the Shapiro-Wilk test, followed by non-parametric Kruskal-Wallis test with Dunn’s multiple-comparison test for *post hoc* group comparisons, using cell numbers or the number of experiments as the basis for ‘n’.

## Acknowledgements

This study was supported by grants from the National Research Foundation of Korea (NRF), funded by the Ministry of Science and ICT (2019R1A2C1086048 to J.W.U; 2017M3C7A1023470 to J.Ko); and a grant from KBSI (T37413 to J.Y.K.).

---

**TABLE S1.**
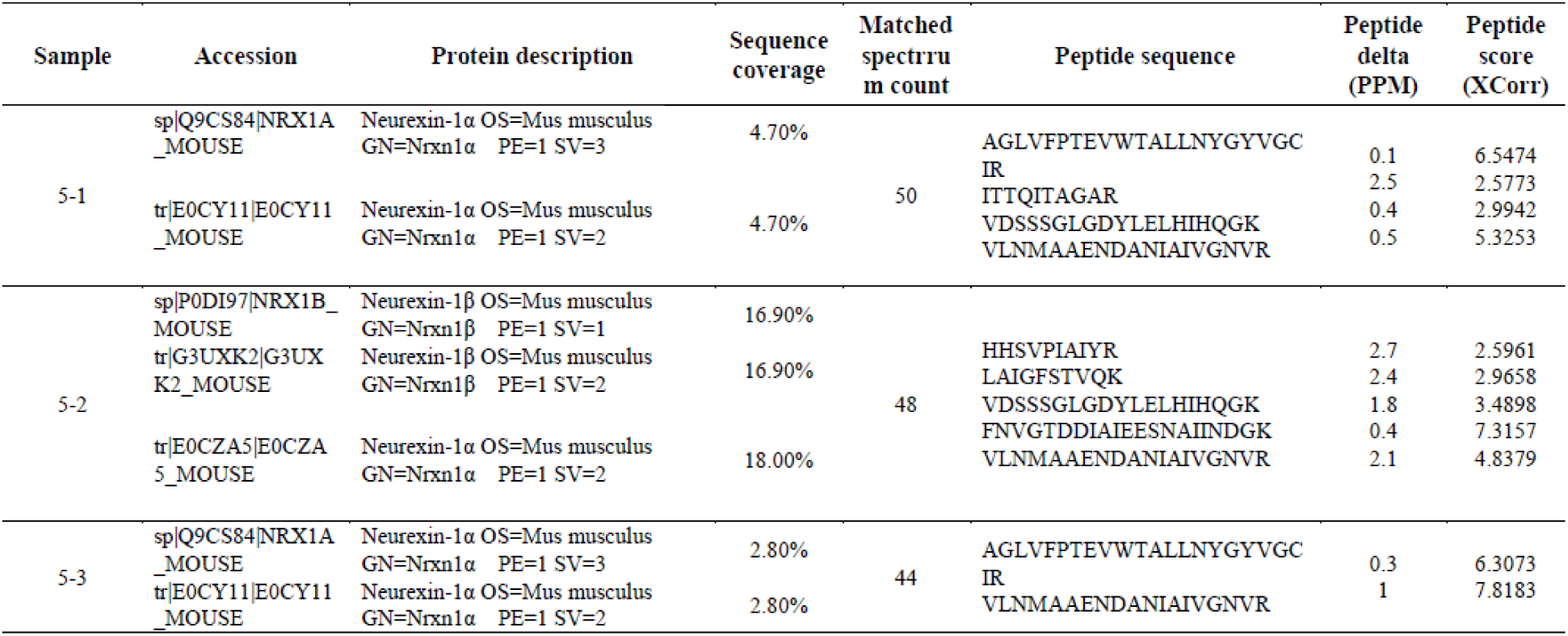
Detailed List of Peptides Identified from Mass Spectrometry

**TABLE S2.** **Offspring Distribution of Different Mouse Genotypes**

|  |  |  |  |  |
| --- | --- | --- | --- | --- |
|  | Nestin-Cre; *Clstn3*<sup>ff</sup> × *Clstn3*<sup>ff</sup> (total : 96) |  |  |  |
|  | ♂ |  | ♀ |  |
| *Clstn3*<sup>ff</sup> |  |  |  |  |
| Nestin-Cre mice | Nestin<sup>+/+</sup> | Nestin<sup>Cre/+</sup> | Nestin<sup>+/+</sup> | Nestin<sup>Cre/+</sup> |
| Observed (%) | 27.08 | 21.88 | 21.88 | 29.17 |
| Expected (%) | 25 | 25 | 25 | 25 |

**Figure 1-Figure supplement 1.**
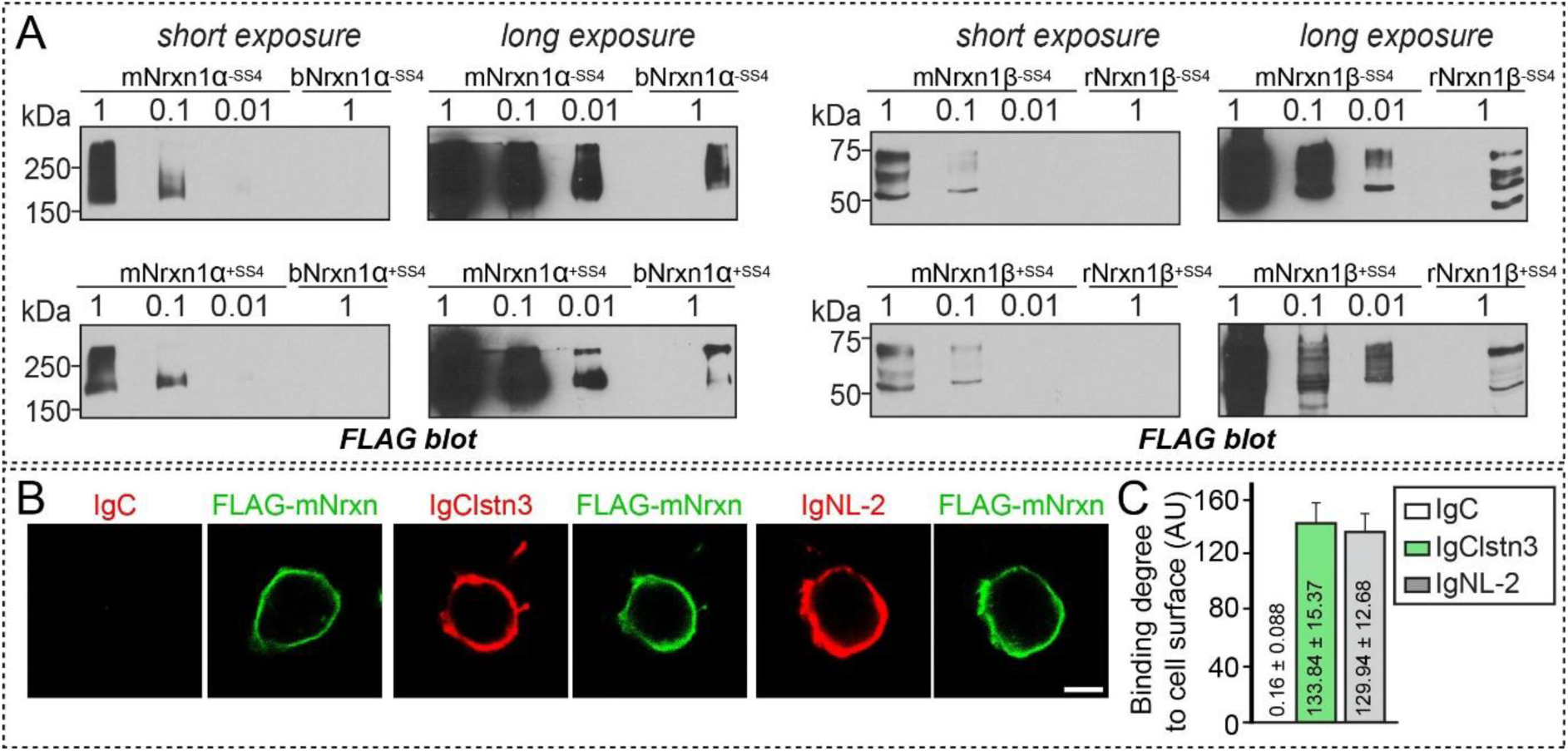
Characterization of Nrxn Constructs Used in The Current Study. **(A)** Immunoblot analyses characterizing the expression levels of Nrxn1α and Nrxn1β constructs. mNrxn1α^+SS4^ and mNrxn1α^-SS4^ encode full-length mouse Nrxn1α with and without an SS4 insert, respectively. bNrxn1α^+SS4^ and bNrxn1α^-SS4^ encode full-length bovine Nrxn1α with or without an SS4 insert, respectively. mNrxn1β^+SS4^ and mNrxn1β ^-SS4^ similarly encode full-length mouse Nrxn1β with or without an SS4 insert, respectively. rNrxn1β^+SS4^ and rNrxn1β^-SS4^ encode full-length rat Nrxn1β with or without an SS4 insert, respectively. mNrxn1α and mNrxn1β constructs were cloned into the pCAGGS vector, whereas bNrxn1α and rNrxn1β constructs, which have long been used to characterize Nrxn-binding ligands, were generated in the Südhof laboratory. Equal amounts of expression plasmids were transfected into HEK293T cells. Serial dilutions of lysates of mNrxn1-expressing cells were required to directly compare expression levels of the indicated Nrxn expression constructs. Note that expression levels of pCAGG mouse Nrxn1 expression plasmids were ∼100-fold higher than those used previously. Molecular mass markers are indicated in kilodaltons (kDa). **(B)** Cell-surface–binding assays. HEK293T cells expressing N-terminally FLAG-tagged mNrxn1β^+SS4^ were incubated with control IgC, IgNL-2, or IgClstn3 fusion proteins and analyzed by immunofluorescence imaging for Ig-fusion proteins (red) and FLAG (green). Scale bars: 10 μm for all images. **(C)** Quantification of cell-surface binding in (**B**). Data are means ± SEMs (number of cells analyzed = 13–15).

**Figure 2-Figure supplement 1.**
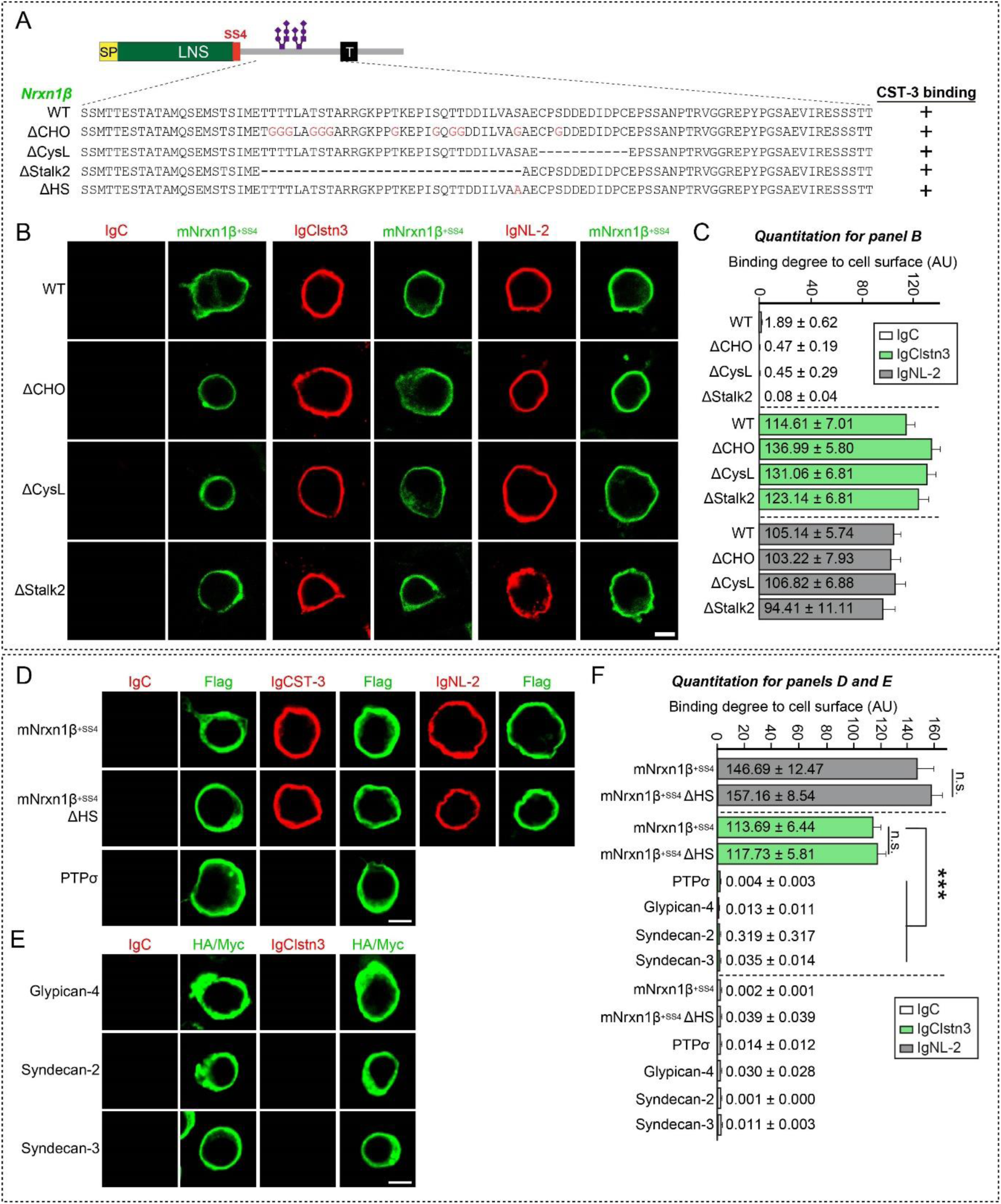
No Effect of Nrxn1β O-glycosylation on Clstn3–Nrxn1β Interactions. **(A)** Diagrams of Nrxn1 constructs used in the cell-surface–labeling assays presented in **(*B*)**. **(B)** Cell-surface–labeling assays. HEK293T cells expressing FLAG-tagged Nrxn1β^+SS4^ WT or its deletion variants (ΔCHO, ΔCysL, or ΔStalk2) were incubated with IgC (control), IgClstn3, or IgNL-2 and analyzed by immunofluorescence imaging for Ig-fusion proteins (red) and FLAG (green). Scale bars: 10 μm for all images. **(C)** Quantification of cell-surface binding in (**B**). Data are means ± SEMs (number of cells analyzed = 11–17). **(D)** Cell-surface–labeling assays. HEK293T cells expressing FLAG-tagged Nrxn1β^+SS4^ WT, its heparan sulfate binding defective mutant (ΔHS), or PTPσ were incubated with IgC (control), IgClstn3, or IgNL-2 and analyzed by immunofluorescence imaging for Ig-fusion proteins (red) and FLAG (green). Scale bars: 10 μm for all images. **(E)** Cell-surface–labeling assays. HEK293T cells expressing HA-tagged Glypican-4, Myc-tagged Syndecan-2 or Syndecan-3 were incubated with IgC (control), or IgClstn3 and analyzed by immunofluorescence imaging for Ig-fusion proteins (red) and HA/Myc (green). Scale bars: 10 μm for all images. **(F)** Quantification of cell-surface binding in (**D**) and (**E**). Data are means ± SEMs (number of cells analyzed = 11–17). (\**p* < 0.05, \*\**p* < 0.01; non-parametric Kruskal-Wallis test with Dunn’s *post hoc* test).

**Figure 2-Figure supplement 2.**
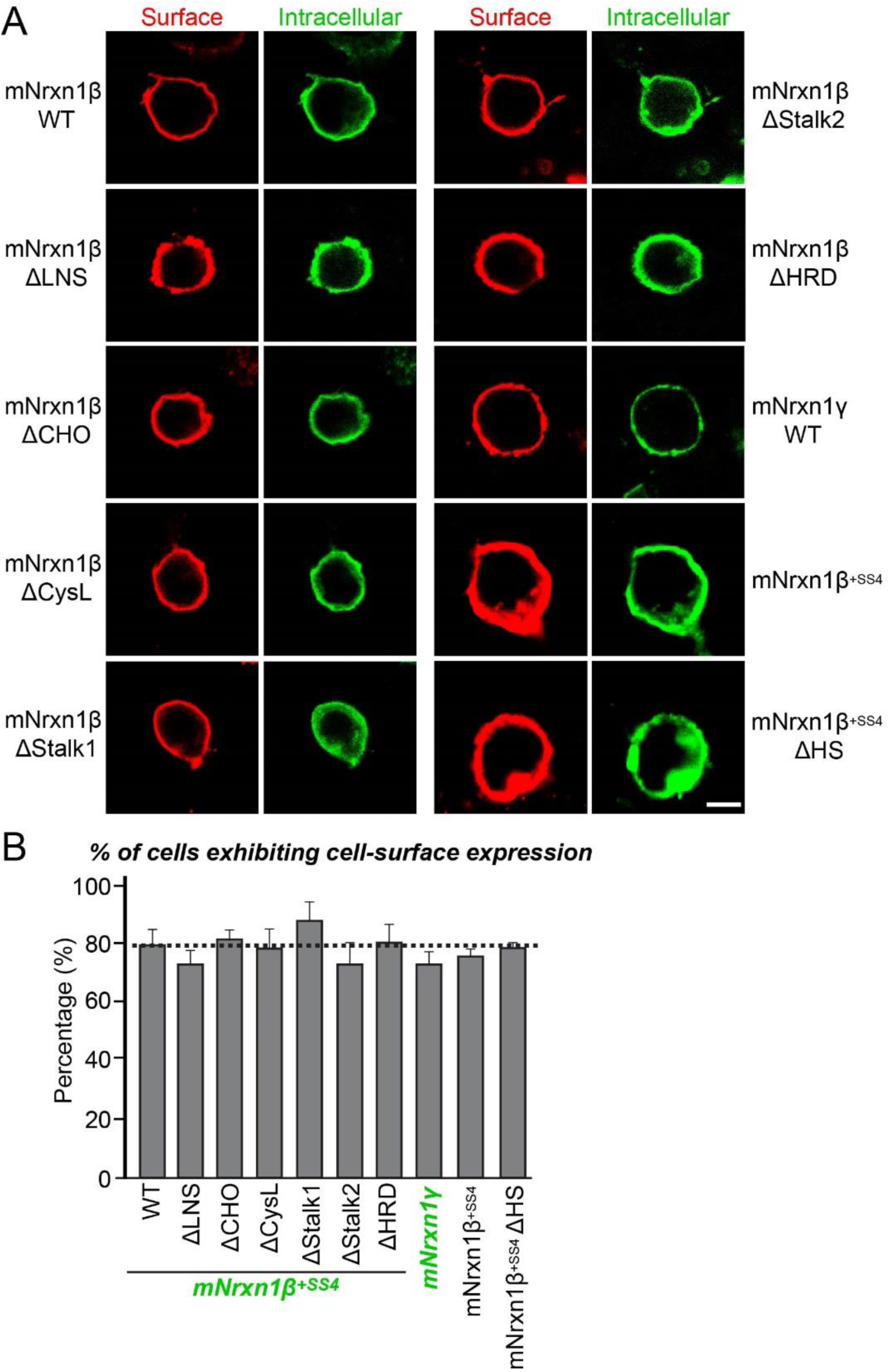
Surface Expression of Nrxn1β Variants Used in the Current Study. **(A)** Surface expression analysis of HEK293T cells expressing FLAG-tagged Nrxn1β^+SS4^ WT, its deletion variants (ΔHRD, ΔLNS, ΔCHO, ΔCysL, ΔStalk1, ΔStalk2 or ΔHS), or FLAG-tagged Nrxn1γ WT. Transfected cells were immunostained with mouse anti-FLAG antibodies and detected with Cy3-conjugated anti-mouse secondary antibodies under non-permeabilized conditions, followed by permeabilization of cells. Cells were then immunostained with rabbit anti-FLAG antibodies and with FITC-conjugated anti-rabbit secondary antibodies. Scale bar: 10 μm for all images. **(B)** Quantification of the proportion of cells exhibiting surface expression of Nrxns. Data are means ± SEMs. ‘n’ denotes the number of cells, as follows: WT mNrxn1β^+SS4^, n = 272; ΔLNS, n = 239; ΔCHO, n = 296; ΔCysL, n = 590; ΔStalk1, n = 240; ΔStalk2, n = 220; ΔHRD, n = 369; mNrxn1γ, n = 161; mNrxn1β^+SS4^, n = 107; and mNrxn1β^+SS4^ ΔHS, n=131.

**Figure 2-Figure supplement 3.**
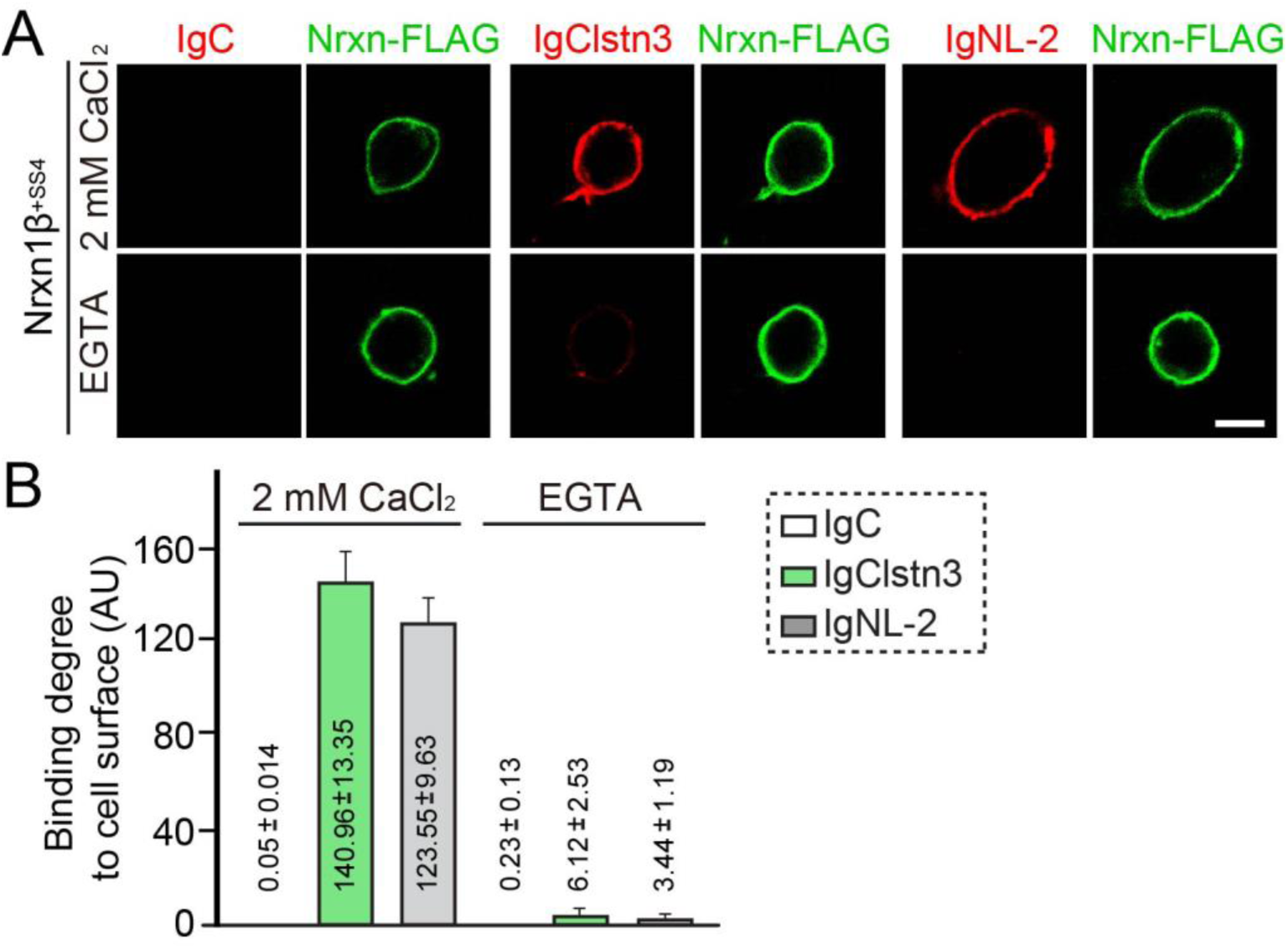
Ca^2+^-dependence of Clstn3–Nrxn1β Interactions. **(A)** Representative images from cell-surface–labeling assays using HEK293T cells transfected with the indicated C-terminally FLAG-tagged Nrxn1β plasmids (Nrxns-FLAG) and incubated with IgC (control Ig-fusion protein), IgNL-2, or IgClstn3. The cells were analyzed by immunofluorescence imaging for Ig-fusion proteins (red) and FLAG (green). All binding reactions were performed in 2 mM CaCl_2_ and 2 mM MgCl_2_, except for the reaction marked “EGTA”, which was performed in the presence of 10 mM EGTA. Scale bars: 10 μm for all images. **(B)** Quantification of cell-surface binding in **(A)**. Data are means ± SEMs (number of cells analyzed = 11–15).

**Figure 3-Figure supplement 1.**
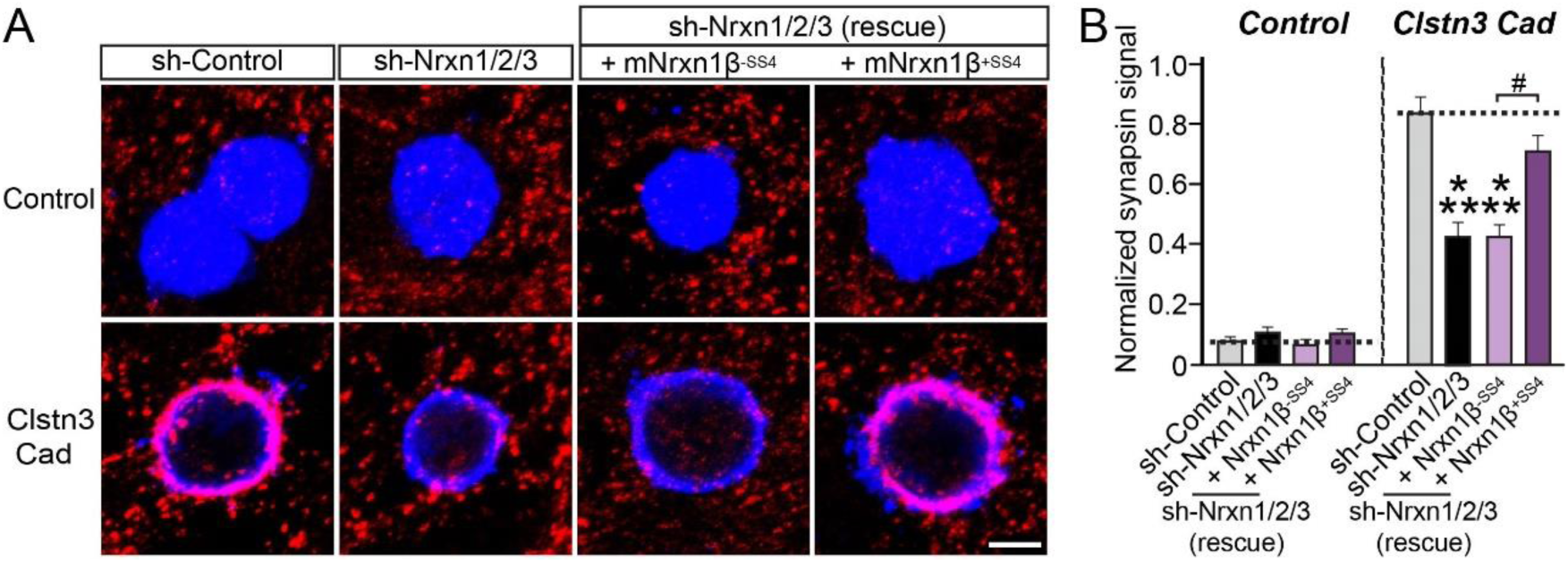
Expression of β-Nrxn Splice Variants Containing an SS4 Insert Rescues the Lost Clstn3 Synaptogenic Activity in Nrxn-deficient Hippocampal Neurons. **(A** and **B)** Effects of Nrxn TKD (sh-Nrxn1/2/3) on the synaptogenic activities of Clstn3 Cad. HEK293T cells expressing EGFP alone (Control) or FLAG-Clstn3 Cad were co-cultured with neurons infected with control lentiviruses (sh-Control) or lentiviruses expressing sh-Nrxn1/2/3, with or without co-expression of shRNA-resistant Nrxn1β splice variants (+mNrxn1β^-SS4^ or +mNrxn1β^+SS4^). Representative images (**A**) of co-cultures immunostained with antibodies to EGFP or FLAG (blue) and synapsin (red). Coincident signals are indicated in violet. Scale bars: 10 μm for all images. Quantification (**B**) of heterologous synapse-formation assays as the ratio of synapsin to EGFP/FLAG fluorescence signals. Dashed lines correspond to baseline control values. Data are means ± SEMs (\*\*\**p* < 0.001; ^#^*p* < 0.05; non-parametric Kruskal-Wallis test with Dunn’s *post hoc* test; ‘n’ denotes the total number of HEK293T cells analyzed).

**Figure 4-Figure supplement 1.**
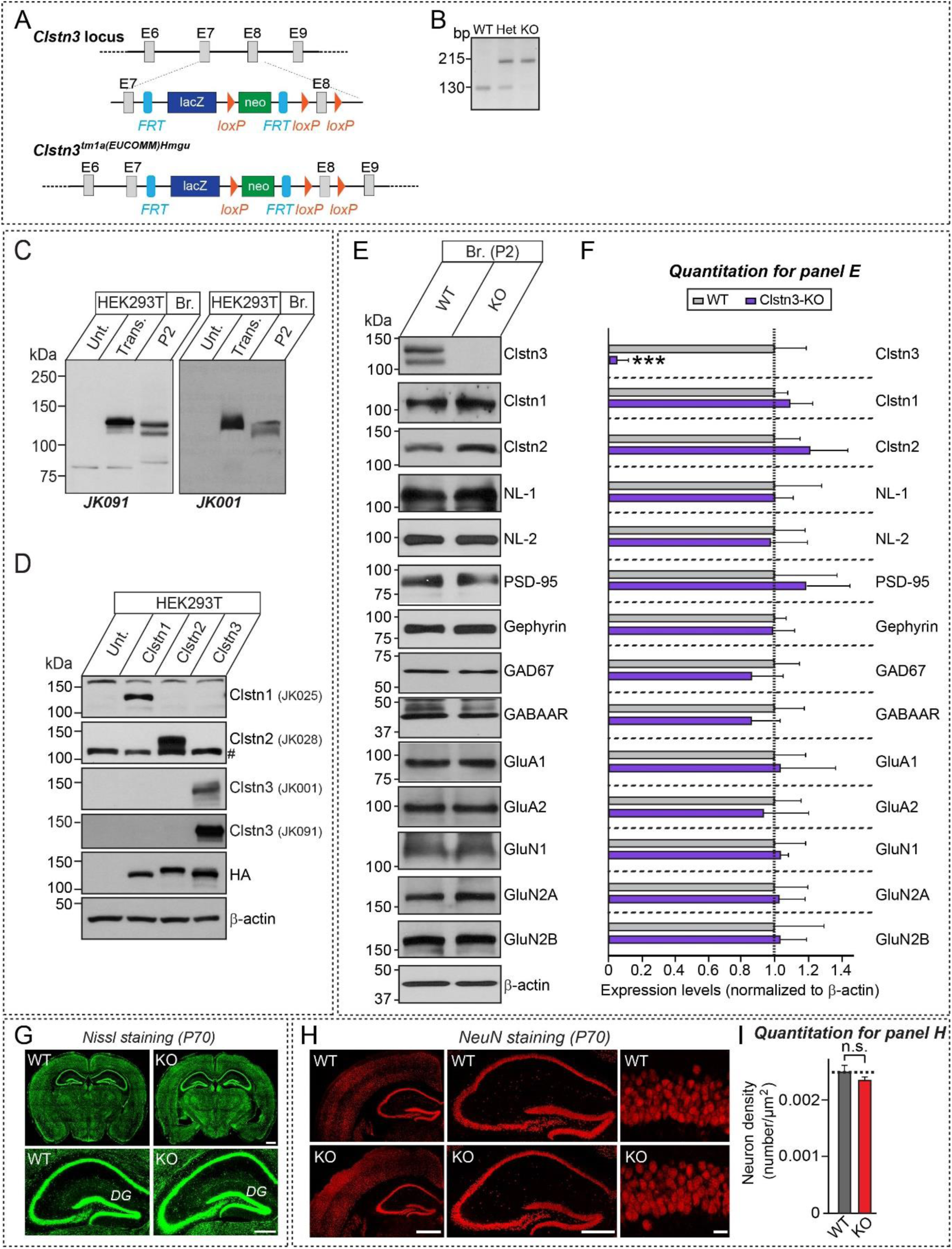
Generation and Characterization of Clstn3^tm1a/tm1a^ Mice. **(A)** Strategy used to generate *Clstn3^tm1a/tm1a^* mice. Arrows flanking the neomycin gene and FLP recombinant target (FRT) or exon 8 (E8) indicate loxP sites. Note that lacZ and neo are two separate markers. **(B)** PCR genotyping of *Clstn3^tm1a/tm1a^* (KO), *Clstn3^tm1a/+^* (Het) and WT mice. **(C)** Immunoblot analyses of Clstn3 antibodies (JK091 and JK001) using lysates from HEK293T cells, untransfected or transfected with HA-tagged Clstn3, and rat brain crude synaptosomal fractions (P2). Br., brain; Unt., untransfected HEK293T cell lysates; Trans., transfected HEK293T cell lysates. **(D)** Cross-reactivity of the Clstn antibodies used in the current study. HEK293T cells, untransfected or transfected with the indicated Clstn3 expression vectors, were immunoblotted with anti-Clstn1 (JK025), anti-Clstn2 (JK028), or anti-Clstn3 (JK091 and JK001) antibodies. The expression of HA-tagged Clstn constructs was confirmed by immunoblotting with anti-HA antibodies. An anti-β-actin antibody was used for normalization. **(E** and **F)** Representative immunoblots (**E**) and summary graphs of synaptic protein levels (**F**) in crude synaptosomal fractions of P42 WT and *Clstn3^tm1a/tm1a^* brains (3–5 pairs), analyzed by semiquantitative immunoblotting. Data are means ± SEMs (\*\*\**p* < 0.001; Student’s *t-*test). **(G)** Normal gross morphology of the *Clstn3^tm1a/tm1a^* brain at P70, as revealed by Nissl staining. Scale bar: 1 mm (top) and 500 μm (bottom). **(H** and **I)** Representative images (**H**) and summary graphs (**I**) of NeuN (neuronal marker) staining showing normal numbers of neurons in hippocampal and cortical regions. Left, forebrain coronal section; middle, hippocampal coronal section; right, hippocampal CA1 stratum pyramidale layer. Scale bar: 1 mm (left), 0.5 mm (middle), and 20 μm (right). Data are means ± SEMs (n = 3 mice each after averaging data from 6 sections/mouse; n.s., not significant).

**Figure 4-Figure supplement 2.**
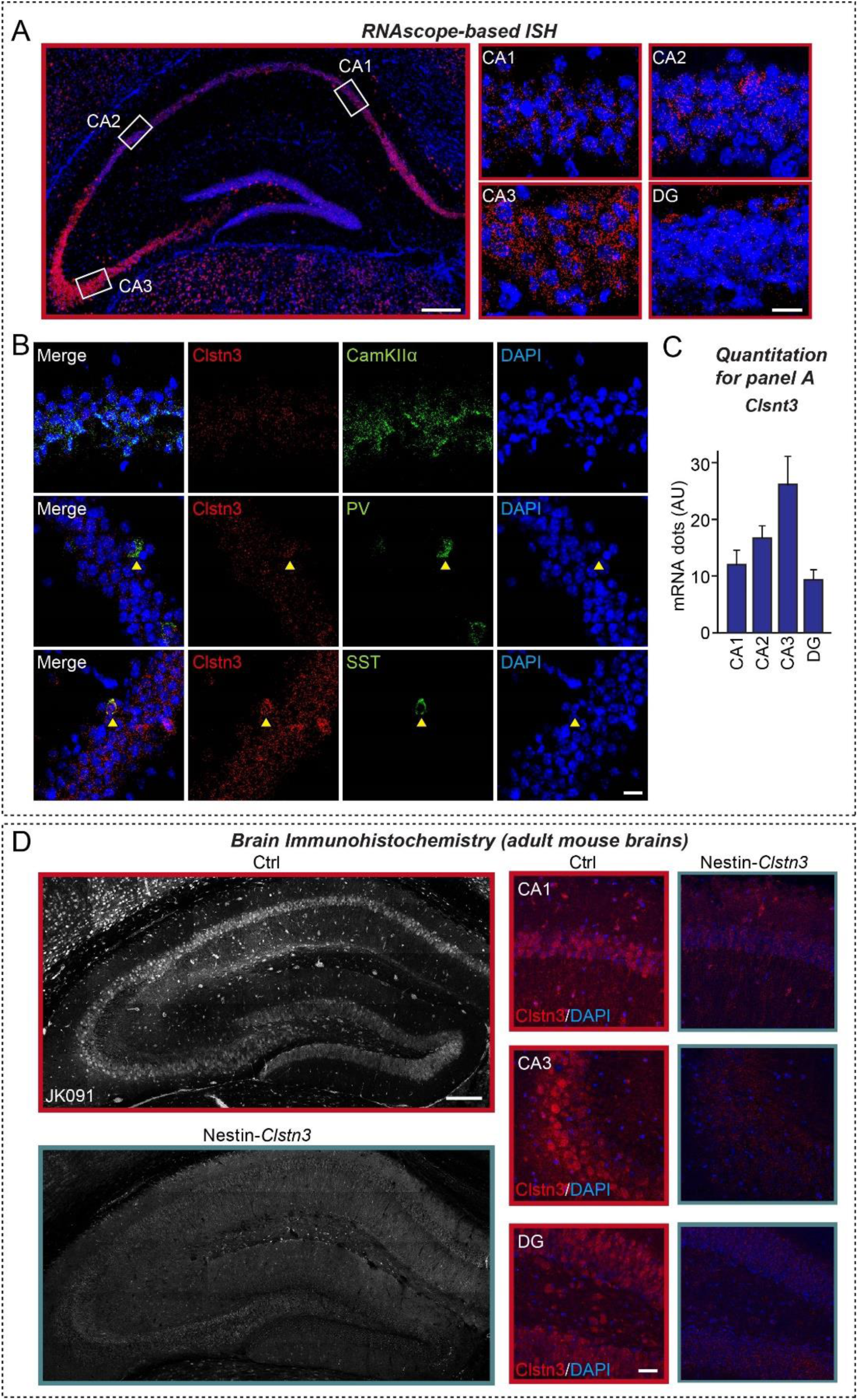
Clstn3 is Expressed in Glutamatergic, PV-positive, and SST-positive Neurons in the Adult Mouse Hippocampus. **(A** and **C)** Expression of *Clstn3* mRNA (red) in various hippocampal regions, as visualized by *in situ* hybridization using RNAscope technology. Nuclei were stained with DAPI (blue). Representative image (**A**) and quantitative analyses (**C**) of RNAscope data are shown. Scale bar: 200 μm for larger color image on the left, and 20 μm for images on the right. (**B**) Expression of *Clstn*3 mRNA (red) in CamKIIα-positive glutamatergic, PV-positive, and SST-positive neurons in hippocampal regions, shown by staining for the indicated gene probes (green) by RNAscope-based *in situ* hybridization, followed by counterstaining with the nuclear dye DAPI (blue). Scale bar: 20 μm. (**D**) Expression of Clstn3 protein in the hippocampus of Clstn3^flox/flox^ (Ctrl) or Nestin-Cre;Clstn3^flox/flox^ (Nestin-*Clstn3*) mice at P42 (6 wks), as visualized by immunohistochemistry using anti-Clstn3 antibodies (JK191). Nuclei were stained with DAPI (blue). Scale bar: 200 μm for larger images on the left, and 20 μm for images on the right.

**Figure 5-Figure supplement 1.**
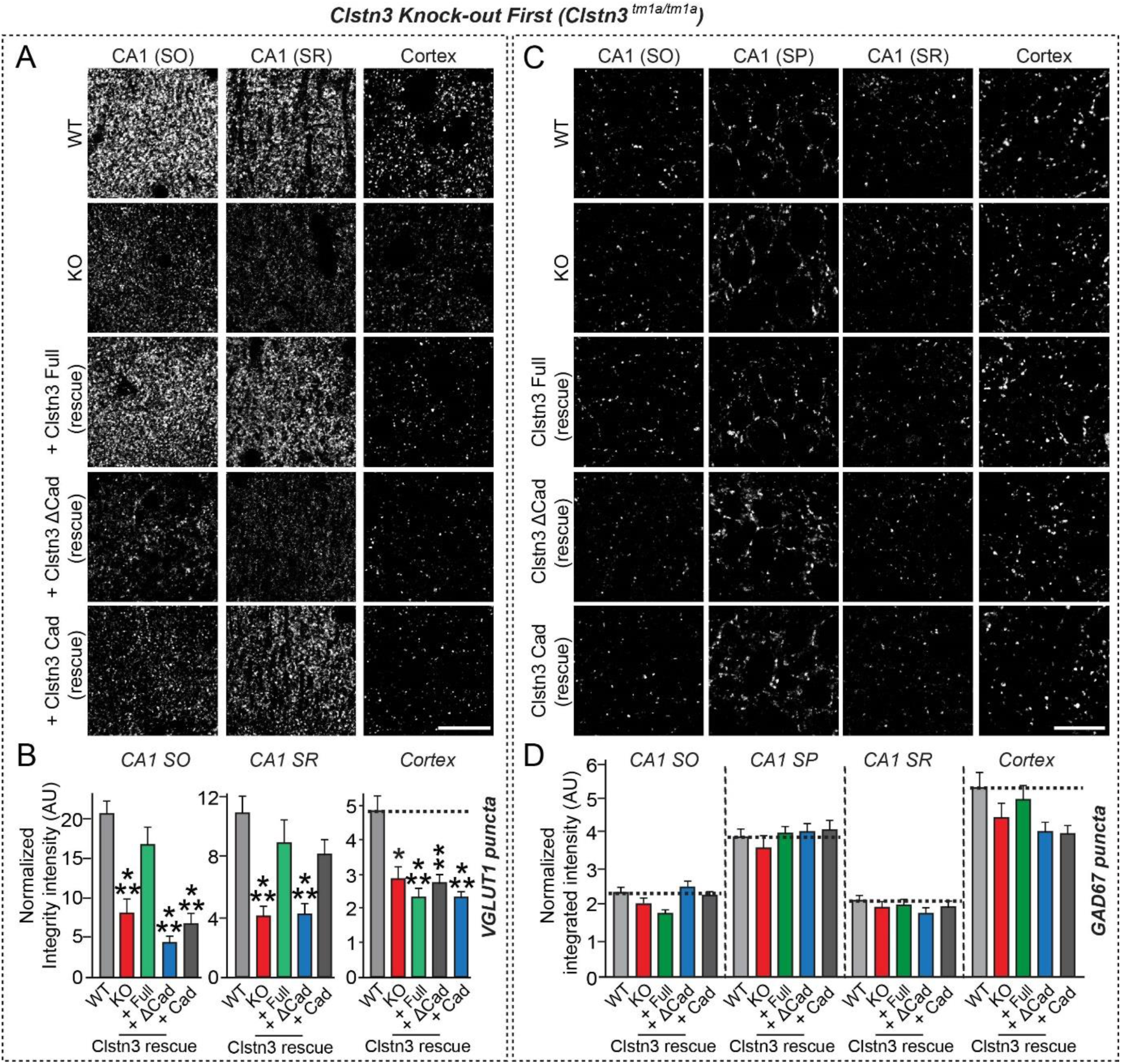
Hippocampal Excitatory Synapse Structural Deficits in Clstn3-KO Mice. **(A)** Representative images of hippocampal CA1 SO and SR regions and cortex 2 wk after stereotactic AAV delivery into WT or *Clstn3^tm1a/tm1a^* mice, followed by immunostaining for the excitatory synapse marker VGLUT1. Scale bar: 20 μm (applies to all images). **(B)** Quantification of the integrated intensity of VGLUT1-positive synaptic puncta. Data are means ± SEMs (\*\*\**p* < 0.001, \*\**p* < 0.01 and \**p* < 0.05; non-parametric Kruskal-Wallis test with Dunn’s *post hoc* test; n = 5 mice each after averaging data from 6 sections/mouse). **(C)** Representative images of the hippocampus CA1 stratum oriens (SO), hippocampus CA1 stratum pyramidale (SP), hippocampus CA1 stratum radiatum (SR), hippocampus dentate gyrus (DG) granulosum, and cortex obtained 2 wk after stereotaxic injection of the indicated AAVs into WT or *Clstn3^tm1a/tm1a^* (KO) mice, followed by immunostaining for the inhibitory synapse marker, GAD67. Scale bar: 20 μm for all images. **(D)** Quantification of the integrated intensity of GAD67-positive synaptic puncta. Data are means ± SEMs (n = 5 mice each after averaging data from 6 sections/mouse).

**Figure 5-Figure supplement 2.**
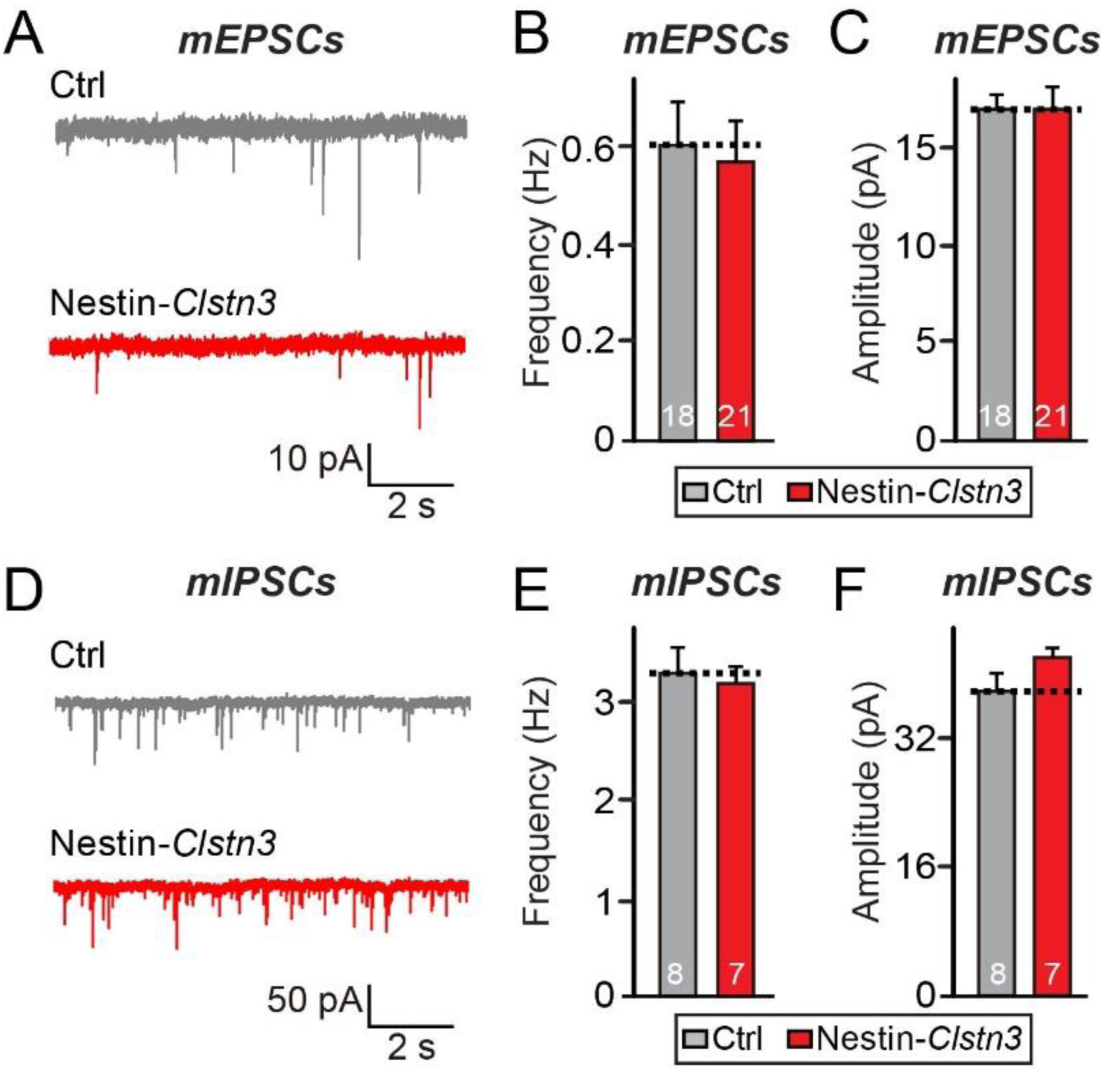
Normal Basal Synaptic Transmission in Clstn3-KO Mice. (**A–C**) Representative traces (**A**) and summary graphs (**B** and **C**) showing the frequency and amplitude of mEPSCs recorded from CA1 pyramidal cells of the hippocampus of *Clstn3*^fl/fl^ (Ctrl) and Nestin-Cre; *Clstn3*^fl/fl^ (Nestin-Clstn3) mice. Graphs show means ± SEMs (Mann-Whitney U test; n denotes the total number of neurons analyzed as follows: Ctrl, n = 18 cells from 8 mice; and Nestin-*Clstn3*, n = 21 cells from 8 mice). (**D–F**) Representative traces (**D**) and summary graphs (**E** and **F**) showing the frequency and amplitude of mIPSCs recorded from CA1 pyramidal cells of the hippocampus of *Clstn3*^fl/fl^ (Ctrl) and Nestin-Cre; *Clstn3*^fl/fl^ (Nestin-*Clstn3*) mice. Graphs show means ± SEMs (Mann-Whitney U test; n denotes the total number of neurons analyzed as follows: Ctrl, n = 8 cells from 2 mice; and Nestin-*Clstn3*, n = 7 cells from 2 mice).

**Figure 7-Figure supplement 1.**
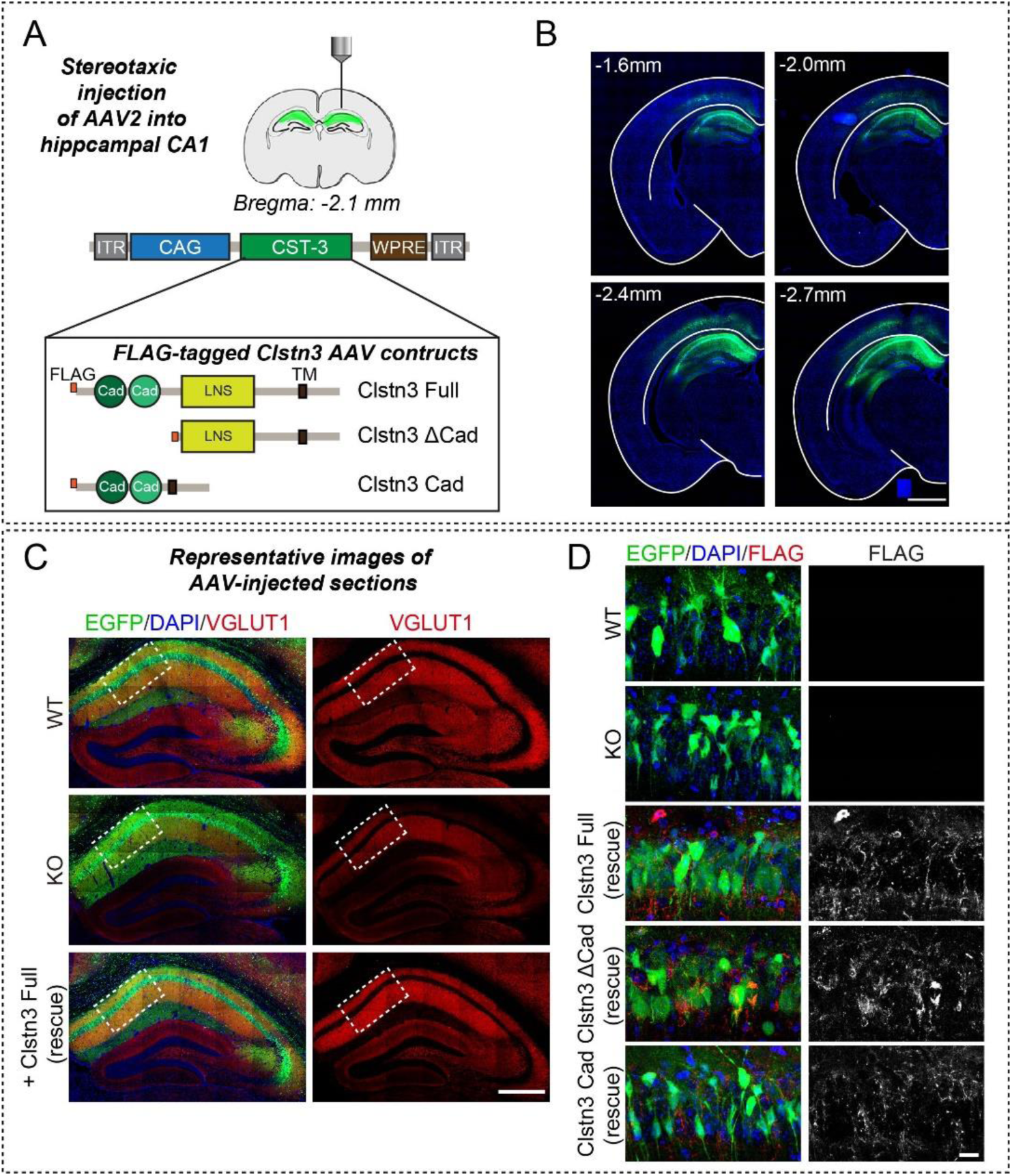
Strategy of AAV-mediated Expression of Clstn3 Proteins in the Hippocampal CA1 Region of Mice. **(A)** Schematic diagram of AAV vectors expressing FLAG-Clstn3 and its deletion variants. AAV viruses expressing the indicated Clstn3 variants were stereotactically co-injected into the hippocampal CA1 region of adult *Clstn3^tm1a/tm1a^* mice with AAV viruses expressing EGFP. **(B)** Representative images illustrating EGFP expression after AAV injection into the hippocampal CA1 region. Different mouse brain coronal sections (-1.6 to -2.7 mm relative to the bregma) were immunostained for EGFP (green) and counterstained with DAPI (blue). Scale bar: 1.0 mm (applies to all images). **(C)** Representative images of AAV-infected neurons in hippocampal CA1 regions, immunostained for EGFP (green) and the excitatory marker VGLUT1 (red), and counterstained with DAPI (blue). Scale bar: 500 μm (for all images). **(D)** Zoomed-in representative images of hippocampal CA1 neurons from Ctrl (WT), Clstn3-KO (KO), and Clstn3 mice infected with the indicated Clstn3 AAV viruses (indicated as ‘rescue’) to confirm AAV-mediated expression of the indicated Clstn3 fragments. Infected mouse brains were immunostained with anti-EGFP (green) and anti-FLAG (red) antibodies, and counterstained with DAPI (blue). Scale bar: 20 μm (applies to all images).

**Figure 7-Figure supplement 2.**
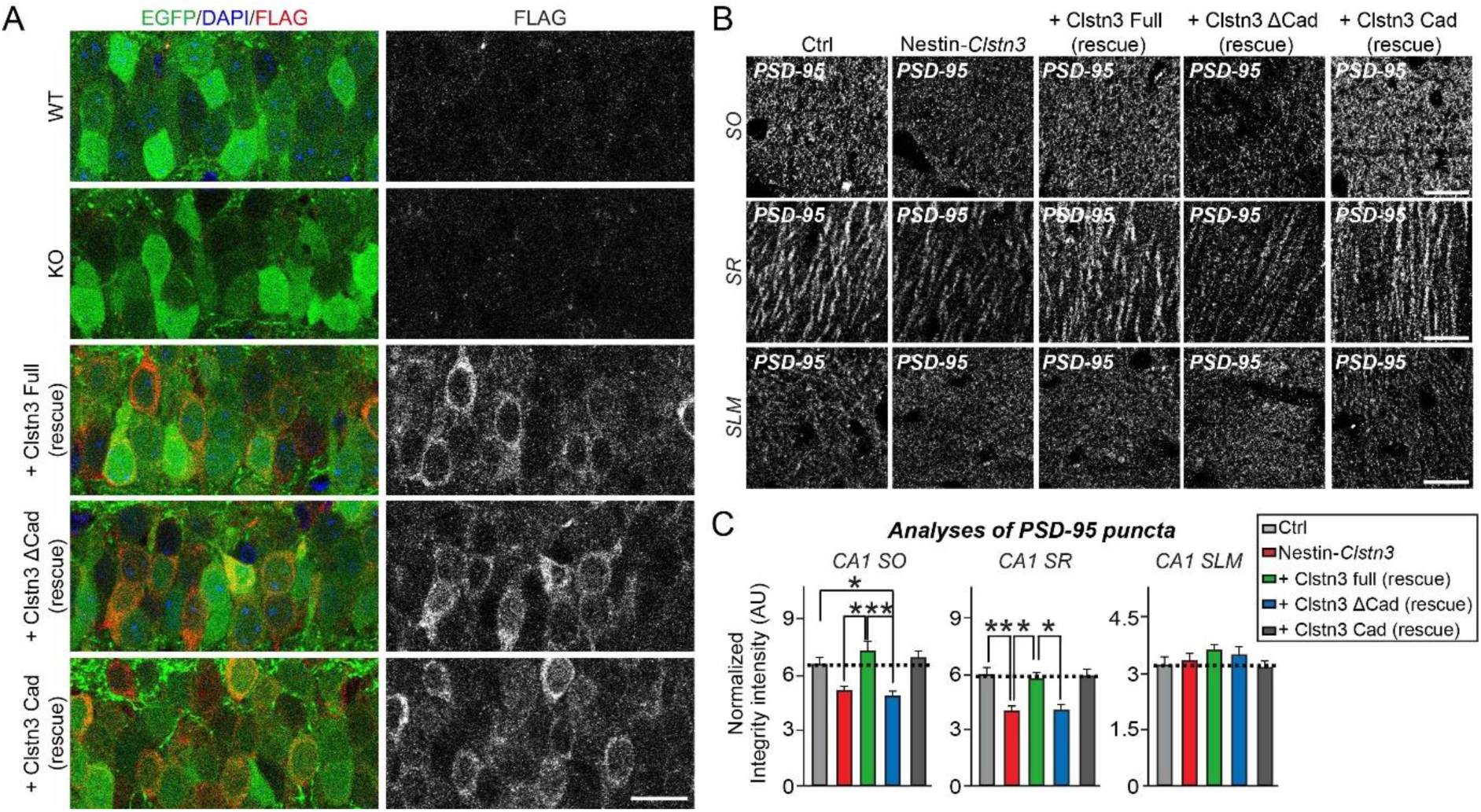
Analysis of Clstn3 Rescue Effects on PSD-95 Puncta in the Hippocampal CA1 Region of Mice. **(A)** Representative images illustrating EGFP and FLAG expression after AAV injection into the hippocampal CA1 region. Injected mouse brain sections were immunostained for EGFP (green), FLAG (red), and counterstained with DAPI (blue). Scale bar: 20 μm (applies to all images). **(B)** Representative images of hippocampal CA1 SO, SR and SLM regions 2 wk after stereotactic injection of the indicated AAVs into *Clstn3^fl/fl^* or *Nestin-Cre;Clstn3^fl/fl^* mice, followed by immunostaining for the excitatory synapse marker PSD-95. Scale bar: 20 μm (applies to all images). **(C)** Quantification of the integrated intensity of PSD-95–positive synaptic puncta. Data are means ± SEMs (\**p* < 0.05 and \*\**p* < 0.01; non-parametric Kruskal-Wallis test with Dunn’s *post hoc* test; n = 5 mice per group after averaging data from 6 sections/mouse).

